# Identification of two pathways mediating protein targeting from ER to lipid droplets

**DOI:** 10.1101/2021.09.14.460330

**Authors:** Jiunn Song, Arda Mizrak, Chia-Wei Lee, Marcelo Cicconet, Zon Weng Lai, Chieh-Han Lu, Stephanie E. Mohr, Robert V. Farese, Tobias C. Walther

## Abstract

Pathways localizing proteins to their sites of action within a cell are essential for eukaryotic cell organization and function. Although mechanisms of protein targeting to many organelles have been defined, little is known about how proteins, such as key metabolic enzymes, target from the ER to cellular lipid droplets (LDs). Here, we identify two distinct pathways for ER-to-LD (ERTOLD) protein targeting: early ERTOLD, occurring during LD formation, and late ERTOLD, targeting mature LDs after their formation. By using systematic, unbiased approaches, we identified specific membrane-fusion machinery, including regulators, a tether, and SNARE proteins, that are required for late ERTOLD targeting. Components of this fusion machinery localize to LD-ER interfaces and appear to be organized at ER exit sites (ERES) to generate ER-LD membrane bridges. We also identified multiple cargoes for early and late ERTOLD. Collectively, our data provide a new model for how proteins target LDs from the ER.

**HIGHLIGHTS:**

- Proteins localize to LDs either during formation or later through ER-LD bridges
- Specific membrane fusion machinery localizes to LDs and mediates protein targeting
- Specific ER exit site proteins associate with LDs and participate in ERTOLD targeting
- Proteomic studies reveal cargoes for early and late ERTOLD targeting

## INTRODUCTION

Compartmentalization of biochemical reactions is a key feature of eukaryotic cells. To enable this compartmentalization, proteins must localize to specific sites in cells, such as the nucleus, mitochondria, or other organelles, where they carry out their functions. Although the protein targeting mechanisms for many organelles is well understood, we know relatively little about how proteins target to the surfaces of lipid droplets (LDs). Compared with other organelles, LDs are unusual since they are bounded by a monolayer of phospholipids that stabilizes their neutral lipid cores (Thiam et al., 2013a). The unusual architecture of LDs presents a challenge for cells: how do cellular proteins target specifically to the LD monolayer surface? Since LDs are ubiquitous organelles that store lipids as metabolic fuel and membrane lipid precursors (Olzmann and Carvalho, 2019; Walther and Farese, 2012; Welte, 2015), this problem is important for understanding the principles of energy storage and also is relevant to human disease. Mutations of LD proteins predispose to or cause metabolic diseases, such as non-alcoholic fatty liver disease/non-alcoholic steatohepatitis (NAFLD/NASH) [e.g., *PNPLA3* (BasuRay et al., 2019; Romeo et al., 2008) and *HSD17B13* (Abul-Husn et al., 2018)] and lipodystrophy [*PLIN1* (Gandotra et al., 2011) and *PCYT1A* (Payne et al., 2014)], and some of the mutations affect localization of the proteins to LDs.

Evolution appears to have solved the problem of targeting proteins to LD surfaces with two principal pathways (Dhiman et al., 2020; Kory et al., 2016). In one, LD proteins are synthesized in the cytoplasm and directly bind LDs. In this <u>cy</u>toplasm <u>to LD</u> (CYTOLD) pathway, most commonly amphipathic helices of soluble proteins directly adsorb and bind to LD surfaces (Pataki et al., 2018; Prévost et al., 2018; Rowe et al., 2016). These protein segments bind preferentially to LDs where packing defects of phospholipids with exposed hydrophobic surfaces are likely more common, larger, and more persistent than in other membranes (Prévost et al., 2018).

The other pathway, ER-to-LD (ERTOLD) targeting, is less well understood. ERTOLD targeting is important for proteins harboring hydrophobic protein segments that are initially inserted into the ER bilayer (Kory et al., 2016; Schrul and Kopito, 2016). The few known cargoes for the ERTOLD pathway include important enzymes of lipid synthesis, such as ACSL3 [long chain acyl-CoA ligase 3] or GPAT4 [glycerol 3-phosphate acyltransferase 4] (Kassan et al., 2013; Wilfling et al., 2013).

How proteins reach LDs from the ER is unclear. Inasmuch as LDs form in the ER, the simplest pathway is to move to the surface of a LD during its formation. Indeed, studies of a small, hydrophobic hairpin sequence derived from GPAT4, known as *LiveDrop,* show that it accumulates on LDs as they are forming in the ER at LD assembly complexes (LDACs), consisting of seipin and accessory proteins (Chung et al., 2019; Wang et al., 2016). The driving force for LD accumulation appears to be a conformational change that allows the protein to adopt an energetically more favorable state on the LD surface than in the ER membrane (Olarte et al., 2020). Similarly, HPos peptide, derived from ACSL3, localizes to LDs during their formation (Kassan et al., 2013).

Surprisingly, full-length GPAT4 does not utilize the same pathway to access LDs. It is excluded from LDs when they form and targets to mature LDs hours later (Wilfling et al., 2013). Light and electron microscopy studies provided evidence that late ERTOLD targeting involves physical continuities—or membrane bridges—between the ER and LDs (Cottier and Schneiter, 2022; Jacquier et al., 2011; Wilfling et al., 2013). How the bridges are formed is unknown. The Arf1/COPI vesicular trafficking machinery (Beck et al., 2009) is required for ERTOLD targeting of proteins, such as GPAT4 (Wilfling et al., 2014) and the major TG lipase ATGL (Beller et al., 2008; Ellong et al., 2011; Soni et al., 2009). Although the function of the Arf1/COPI machinery in this process is uncertain, it may promote membrane bridges forming between ER and LDs (Thiam et al., 2013b; Wilfling et al., 2014).

Here we sought to uncover the pathways of ERTOLD protein targeting. Specifically, we tested the hypothesis that there are early and late pathways of ERTOLD targeting, and we investigated the mechanism behind the formation of ER-LD membrane bridges that mediate late ERTOLD targeting. Using an unbiased screening approach, we identified the protein machinery for late ERTOLD targeting that supports heterotypic organelle fusion of the ER and LDs. This trafficking appears to occur at ERES, a subdomain of the ER important for protein export. Capitalizing on these insights, we systematically identified cargoes for the late ERTOLD targeting pathway.

## RESULTS

### ERTOLD cargoes access LDs at different time points during LD formation

*LiveDrop,* but not full-length GPAT4, accesses LDs during their formation (Wang et al., 2016; Wilfling et al., 2013). To test whether ERTOLD targeting during LD formation is specific to *LiveDrop* or applies to full-length native proteins, we co-expressed fluorescently tagged GPAT4 and LD-associated hydrolase (LDAH), another known ERTOLD cargo (Thiel et al., 2013), in *Drosophila* cells. LDAH is a serine hydrolase implicated in the turnover of cholesterol esters in human macrophages and lipid storage in *Drosophila* (Goo et al., 2014; Kory et al., 2017; Thiel et al., 2013). LDAH is enriched on LDs as early as 30 min after induction of LD formation (by incubating cells in oleate-containing medium), but GPAT4 is enriched on LDs ∼3 hours later (**Figure 1A**).

**Figure 1.**
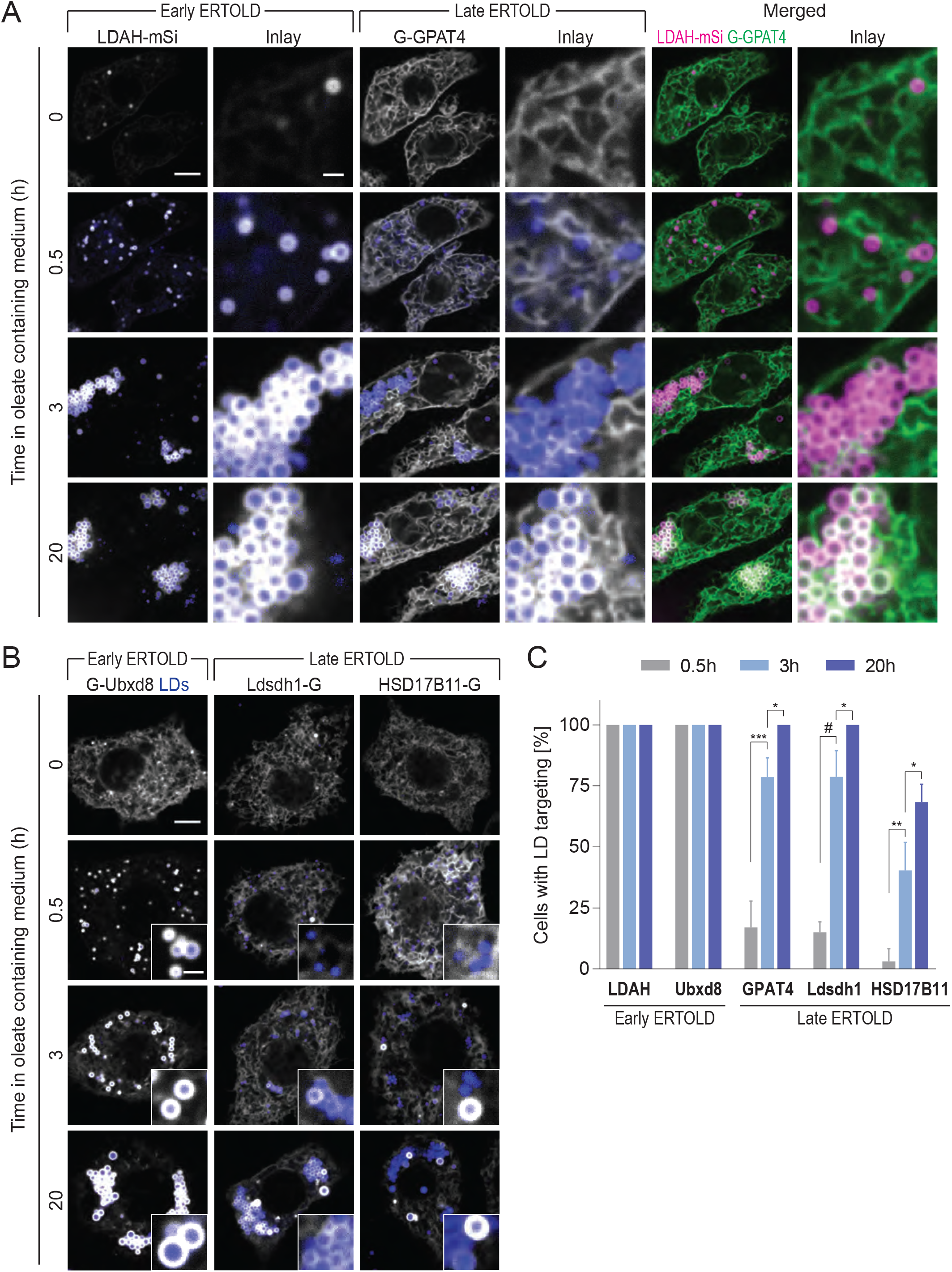
ER proteins target LDs early (early ERTOLD) or late (late ERTOLD) after LD induction. (A) ER proteins LDAH and GPAT4 target LDs early or late, respectively, upon LD biogenesis. Confocal imaging of live *Drosophila* S2 R+ cells stably overexpressing EGFP-GPAT4 transfected with a LDAH-mScarlet-I(mSi) encoding construct at given timepoints after 1 mM oleic acid treatment. LDs were stained with LipidTOX Deep Red Neutral Lipid Stain. Scale bar, 5 and 1 μm (inlay). (B) Ubxd8 targets LDs early, and Ldsdh1 and HSD17B11 target LDs late upon LD biogenesis. Confocal imaging of live wildtype cells transiently transfected with EGFP tagged constructs at given timepoints after 1 mM oleic acid treatment. LDs were stained with monodansylpentane (MDH). Representative images are shown. Scale bar, 5 and 1 μm (inlay). (C) Bar graph showing percentage of cells with LD targeting over time from the imaging experiment in (B), except LDAH-EGFP and EGFP-GPAT4 (not shown). For HSD17B11, cells with LD targeting were defined as those with >2 LDs with protein targeting in the imaging plane (see **Figures S1A&B**). Mean ± SD, n = 3 independent experiments (10–16 cells each). One-way ANOVA, *p<0.05, **p<0.01, ***p<0.001, #p<0.0001.

We also tested the targeting kinetics of several other LD proteins with hydrophobic regions that localize to the ER in the absence of LDs (Liu et al., 2018; Olzmann et al., 2013; Thul et al., 2017; Yu et al., 2018). Ubxd8, a recruitment factor for the p97 segregase, behaved comparable to LDAH and targeted LDs during formation, whereas the enzymes Ldsdh1 and HSD17B11 localized to LDs at later times, similar to GPAT4 (**Figures 1B and 1C**). Notably, overexpressed HSD17B11 targeted only some LDs, suggesting additional determinants of LD targeting for this protein (**Figures S1A and S1B**). Taken together, LD proteins appear to use different ERTOLD pathways to access LDs: some target during formation, and others later in LD biogenesis.

### A genome-wide screen identifies mediators of late ERTOLD targeting

The inability of some ERTOLD proteins to access LDs during their formation raises the question of how they target mature LDs at later times. To address this, we systematically screened the genome for factors required for GPAT4 targeting to LDs (**Figure 2A**). Specifically, we determined the effects of RNAi-mediated protein depletions on LD targeting of stably expressed, fluorescently tagged GPAT4. Duplicate experiments were performed for the entire genome, collecting eight images for each knockdown and generating ∼1.2 million images in total. Automated image analysis segmented nuclei and LDs to quantify the level of GPAT4 at LDs for each cell (**Figure S1C**). We expressed the results as a LD targeting ratio, calculated by dividing the fluorescent signal of GPAT4 on LDs by the signal outside the LDs, and the median across all cells for each knockdown was reported as the final readout. Plotting the distribution of LD targeting ratios across all gene knockdowns revealed a normal distribution around a median value of 2.42, similar to the result for negative control RNAi against *LacZ* (not expressed in *Drosophila* cells; **Figures 2B and 2C**). As expected, depleting most gene products had no effect on GPAT4 targeting to LDs. In contrast, depleting the positive control proteins βCOP or Arf1 (Wilfling et al., 2014) greatly decreased LD targeting of GPAT4, whereas depleting seipin increased targeting (Wang et al., 2016) (**Figures 2B and S1D**). These controls were reproducible, and results from replicate screens correlated well (R=0.7645; **Figure S1E**). To make the results more accessible and allow efficient data mining, we will deposit all images and analyses at the Lipid Droplet Knowledge Portal [http://lipiddroplet.org/; (Mejhert et al., 2021)].

**Figure 2.**
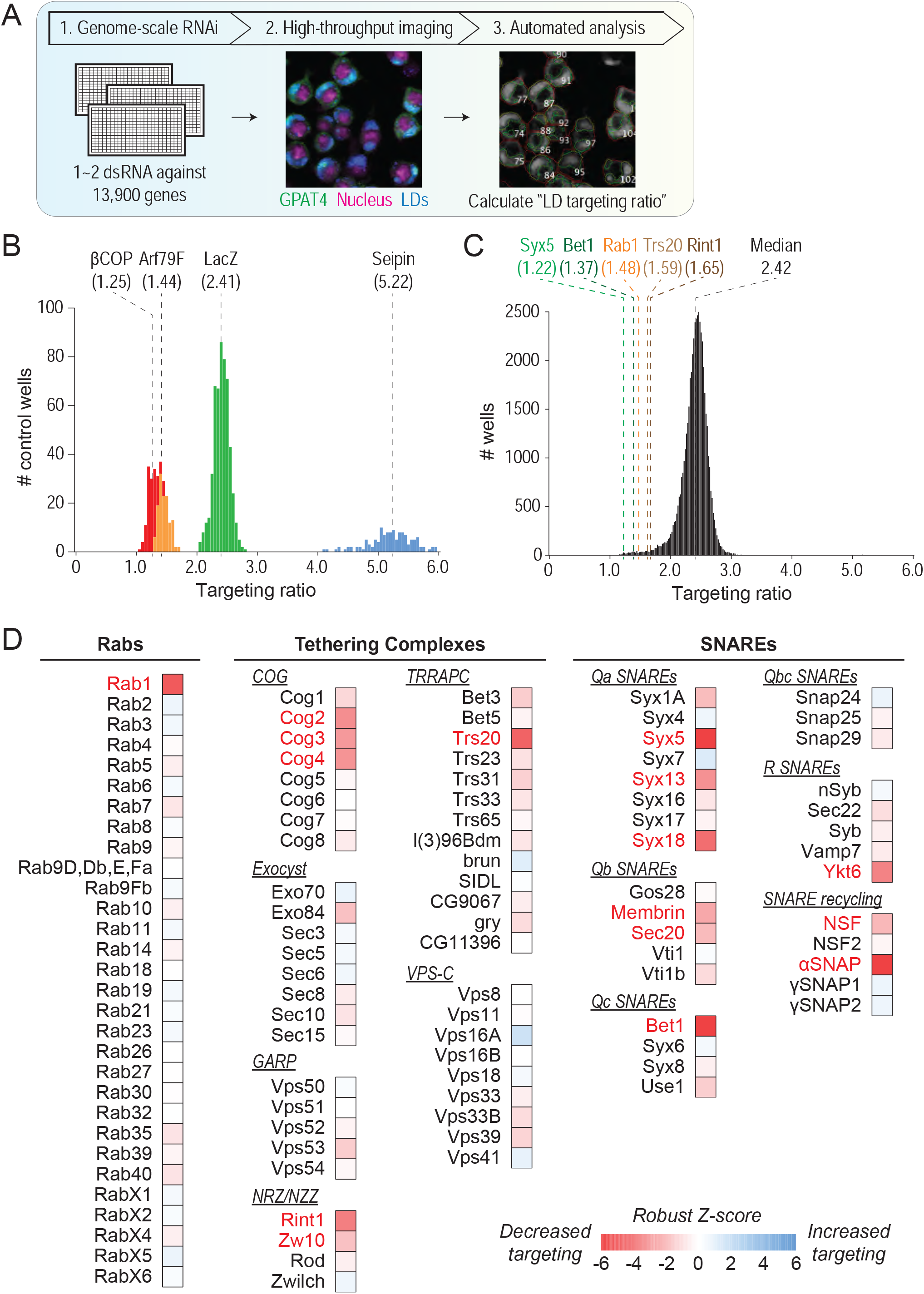
Genome-scale imaging screen reveals that the membrane-fusion machinery is required for GPAT4 targeting to LDs. (A) Overview of genome-scale imaging screen (B) Histogram of targeting ratios of screen controls (n = 528 for LacZ; n = 132 for βCOP, Arf79F, and Seipin). Dotted lines indicate median values for each control. (C) Histogram of all targeting ratios in the screen (n = 50,688). Median of all targeting ratios is indicated in black. Targeting ratios of select screen hits are indicated in different colors. (D) Heatmap of robust Z-scores for different classes of membrane fusion machinery (Rabs, tethering complexes, and SNAREs) from the imaging screen. Genes of which knockdown results in robust Z-score < -2.5 are highlighted in red.

As a conservative cut-off for identifying screen hits, we focused on genes with a robust z-score of less than -2.5 or greater than 2.5, indicating that the LD targeting ratio was at least 2.5 median absolute deviations from the overall median. These cutoffs yielded 896 genes that decreased and 214 genes that increased GPAT4 targeting upon knockdown (of ∼13,900 genes tested), excluding ribosomal, proteasomal, or spliceosomal genes. Analysis of the genes that reduced targeting revealed many genes encoding membrane-trafficking proteins. This was corroborated by gene ontology analysis (Eden et al., 2009) that showed enrichment of genes involved in Golgi apparatus vesicular transport (p=6.77E-08) and vesicle-mediated transport (p=3.59E-06), as well as by enrichment of protein complexes (Vinayagam et al., 2013) involved in vesicle fusion and tethering (**Figure S1F**).

### Specific membrane-fusion factors are required for late ERTOLD targeting

Among the genes required for GPAT4 targeting, we detected a Rab protein, a membrane tether, specific SNAREs, and auxiliary proteins that recycle the membrane-fusion machinery (**Table S1** lists the gene nomenclature in this study).

Of the 30 *Drosophila* Rab GTPases, only Rab1 (robust Z-score = -5.5) was required for GPAT4 targeting to LDs in the screen (**Figure 2D**). To validate the specificity of this finding and the requirement of Rab1 in late ERTOLD targeting, we designed 2–3 additional dsRNAs against Rab proteins implicated in LD biology (Schroeder et al., 2015; Tan et al., 2013; Wang et al., 2012; Wu et al., 2014; Xu et al., 2018) and tested whether they are required for LD localization of GPAT4, fluorescently tagged at its endogenous genomic locus (**Figures 3A and S2A**). In agreement with the screen result, Rab1 depletion abolished GPAT4 targeting to LDs. In contrast, depleting Rab7, Rab8, Rab18, Rab32, or Rab40 had no effect on GPAT4 targeting. Expressing tagged wildtype Rab1 in cells depleted of Rab1 (with dsRNA against 5′ UTR) rescued GPAT4 targeting to LDs, indicating that the effect is specific to the RNAi-mediated depletion of Rab1 (**Figures S2B and S2C**). Expression of the Rab1 N124I mutant, which acts as a dominant-negative to endogenous Rab1 by sequestering its activating guanine nucleotide exchange factor [GEF; (Satoh et al., 1997)], impaired endogenous GPAT4 targeting to LDs (**Figures S2B and S2C**).

**Figure 3.**
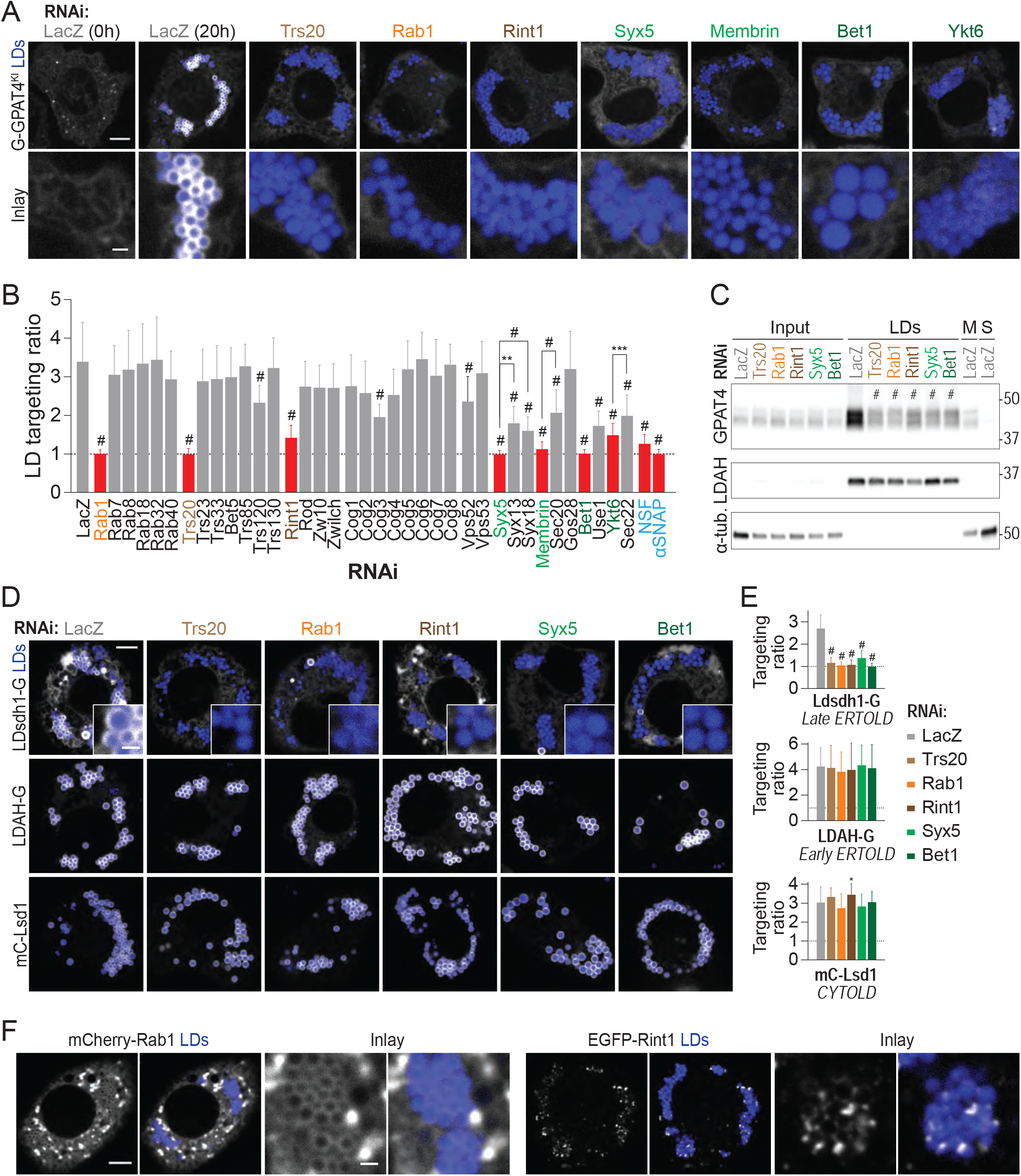
Membrane-fusion regulator, tether, and SNAREs are required for late ERTOLD targeting. (A) Depletion of specific Rab, membrane-tethering complex components, and SNAREs abolishes endogenous GPAT4 targeting to LDs. Confocal imaging of live EGFP-GPAT4 endogenous knock-in cells upon RNAi of membrane-fusion machinery components, followed by a 20-hr incubation in oleate-containing medium (except for 0 h timepoint for LacZ RNAi). LDs were stained with MDH. Representative images are shown. Scale bar, 5 and 1 μm (inlay). (B) Bar graph showing targeting ratios from the imaging experiment in (A) and **Figure S2A**. Mean ± SD, n = 18–45 cells from 2–3 independent experiments. Red color highlights knockdowns that abolish GPAT4 targeting to LDs on imaging. One-way ANOVA, **p<0.01, ***p<0.001, #p<0.0001, compared to LacZ unless otherwise indicated. (C) Depletion of specific Rab, membrane-tethering complex components, and SNAREs reduces GPAT4 amount in LD fractions. Western blot analysis of fractions of wildtype cells upon RNAi, followed by a 20-hr incubation in oleate-containing medium. Indicated on the left-hand side of the blot are protein targets of the primary antibodies, and on the right-hand side are ladder positions. Input = whole-cell lysate, M = membranes, S = soluble fraction. Intensities of GPAT4 bands in LD fractions relative to LacZ control (1.00) were: Trs20^#^ (0.34 ± 0.06), Rab1^#^ (0.28 ± 0.03), Rint1^#^ (0.37 ± 0.04), Syx5^#^ (0.35 ± 0.03), and Bet1^#^ (0.38 ± 0.04) (mean ± SD, n = 3 independent experiments). One-way ANOVA, #p<0.0001, compared to LacZ. (D) Depletion of specific Rab, membrane-tethering complex components, and SNAREs impairs targeting of Ldsdh1 (late ERTOLD) but not of LDAH (early ERTOLD) or Lsd1 (CYTOLD). Confocal imaging of live wildtype cells upon RNAi of fusion machinery, followed by transient transfection of EGFP tagged constructs and a 20-hr incubation in oleate-containing medium. Scale bar, 5 and 1 μm (inlay). (E) Bar graph showing targeting ratios from the imaging experiment in (D). Mean ± SD, n = 35– 50 cells from 3 independent experiments. One-way ANOVA, *p<0.05, #p<0.0001, compared to LacZ. (F) Localization of transiently transfected mCherry-Rab1 or EGFP-Rint1 with respect to LDs. Scale bar, 5 and 1 μm (inlay).

Among membrane tethering complex components, GPAT4 targeting to LDs was reduced upon depletion of Cog2 (robust Z-score = -3.5), Cog3 (-3.2), Cog4 (-3.3), Rint1 (-4.1), Zw10 (- 2.8), and Trs20 (-4.9) (**Figure 2D**). In follow-up studies with additional dsRNA designs, depletion of Cog2, Cog3, and Cog4 led to much smaller LDs but did not impair LD targeting of GPAT4 (**Figure S2A**). We suspect that the apparent low targeting ratios may have resulted from small LDs and the limited resolution of the automated confocal microscope. In contrast, depletion of Trs20 and Rint1 abolished LD targeting of tagged endogenous GPAT4 and resulted in LD targeting ratios close to 1 (**Figures 3A and 3B**). Yet, loss of the other components of TRAPP complexes (e.g., Trs23, Trs33, Bet5, Trs85, Trs120, or Trs130) or NRZ/NZZ complexes (e.g., Rod, Zw10, Zwilch) did not impair LD targeting of endogenous GPAT4 (**Figures 3B and S2A**). Depleting Vps52 and Vps53 (components of GARP complex) did not affect GPAT4 targeting to LDs (**Figures 3B and S2A**). Thus, of the membrane tethering factors, Trs20 and Rint1 were required for GPAT4 targeting.

In vesicular fusion, four SNARE proteins (one from each of Qa, Qb, Qc, and R classes) localize to the vesicle and target membrane and assemble to fuse the compartments (Hong, 2005; Söllner et al., 1993). For repeated cycles, the auxiliary proteins NSF and αSNAP disassemble the post-fusion SNARE complex (Südhof and Rothman, 2009; Ungar and Hughson, 2003; Whiteheart et al., 1994; Zhao et al., 2015). In the screen, each known *Drosophila* SNARE protein was individually depleted. Depleting several Qa SNAREs (Syx5, robust Z-score = -6.7; Syx13, -3.4; Syx18, -4.5) and Qb SNAREs (membrin, -2.7; Sec20, -2.9) and a single Qc SNARE Bet1 (-6.1) and R SNARE Ykt6 (-3.9) reduced GPAT4 targeting to LDs (**Figure 2D**). In experiments with additional dsRNAs and endogenous GPAT4 knock-in cells, depletion of candidates reduced but did not abolish LD targeting of GPAT4 *per se*, except for a single SNARE of each class. Specifically, depletion of Syx5 (Qa), membrin (Qb), Bet1 (Qc), or Ykt6 (R) abolished GPAT4 targeting (**Figures 3A, 3B, and S2A**). Depleting NSF (robust Z-score = -3.3) or αSNAP (-6.4), but not NSF2, γSNAP1, or γSNAP2, reduced GPAT4 targeting to LDs in the screen, and these results were confirmed with additional dsRNAs in cells expressing endogenously tagged GPAT4 (**Figures 3B and S2A**). Finally, expressing the Syx5 1-445 truncation mutant (missing the C-terminal transmembrane segment) or the NSF-E329Q mutant (defective in ATP hydrolysis), which act as dominant-negatives to endogenous Syx5 and NSF, respectively (Dascher et al., 1994; Whiteheart et al., 1994), impaired LD targeting of endogenously tagged GPAT4 (**Figures S2B and S2C**).

We also examined the targeting of endogenous GPAT4 by immunoblotting lysate fractions from cells depleted of Trs20, Rab1, Rint1, Syx5 or Bet1. Endogenous GPAT4 amounts were decreased in LD fractions and increased in microsomal fractions for each of these factor knockdowns (**Figures 3C and S3A**).

Importantly, depletion of membrane fusion machinery specifically affected ERTOLD cargo, as depleting Trs20, Rab1, Rint1, Syx5, or Bet1 did not affect the LD delivery of CYTOLD cargoes, such as Lsd1, CGI-58, or CCT1 (**Figures 3D-E and S3F-G**). The targeting phenotype was further specific to the late ERTOLD pathway, as depletion of Trs20, Rab1, Rint1, Syx5, or Bet1 impaired LD targeting of Ldsdh1 (**Figures 3D and 3E**) and HSD17B11 (**Figures S3B-E**) but not of the early ERTOLD cargoes LDAH or Ubxd8 (**Figures 3C-E and S3F-G**).

### Membrane-fusion factors Rab1 and Rint1 localize to LDs

To determine if the membrane-fusion machinery identified by the screen acts directly at LDs, we analyzed the localization of these factors in cells. Review of published data (Krahmer et al., 2018) supported localization of specific SNARE proteins at LDs; proteomic analysis of liver LDs showed that three of the four SNARE orthologs required for late ERTOLD in *Drosophila* cells (i.e., Stx5 (ortholog of Syx5), Bet1l (ortholog of bet1), and Ykt6) were enriched in LD fractions (**Figure S4C**). However, the site of SNARE protein action is difficult to analyze because these proteins act transiently in numerous membrane-fusion reactions. We therefore focused on the upstream factors, Rab1 and Rint1. To analyze their localizations, we expressed tagged versions of Rab1 or Rint1 in *Drosophila* cells, induced LDs by growing cells for 20 hours in oleate-containing medium, and analyzed their distribution by confocal microscopy. In agreement with prior reports of Rab1 enrichment in LD proteomes (Bersuker et al., 2018; Krahmer et al., 2013), mCherry-Rab1 formed ring-like intensities around LDs (**Figure 3F**). In comparison, EGFP-Rint1 showed a punctate localization with most of the signals adjacent to or overlapping with LDs (**Figure 3F**). Trs20 localized to LDs in a small fraction of cells when overexpressed (**Figure S4A**). When co-expressed with Rint1, both robustly localize to LDs, suggesting Trs20 is recruited to LDs by Rint1 (**Figure S4B**).

### ER exit sites are required for late ERTOLD protein targeting

Membrane fusion in cells is often spatially organized to specific domains of organelles. It was thus noteworthy that a second category of membrane-trafficking factors required for GPAT4 targeting from the screen was genes involved in ERES organization and function (**Figure S1F**). ERES are special ER domains that form transport carriers with protein cargoes destined for secretion via the Golgi apparatus (Bannykh et al., 1996). Our screen identified most proteins that function at ERES as required for GPAT4 targeting to LDs, including Sec12, Sec16, Tango1, and most of COPII coat components (Sar1, Sec23, Sec24AB, Sec24CD, and Sec13) (**Figure 4A**). In contrast, other proteins implicated in secretory trafficking (e.g., coiled-coil tethering proteins) were not required for GPAT4 targeting to LDs (**Figure 4A**).

**Figure 4.**
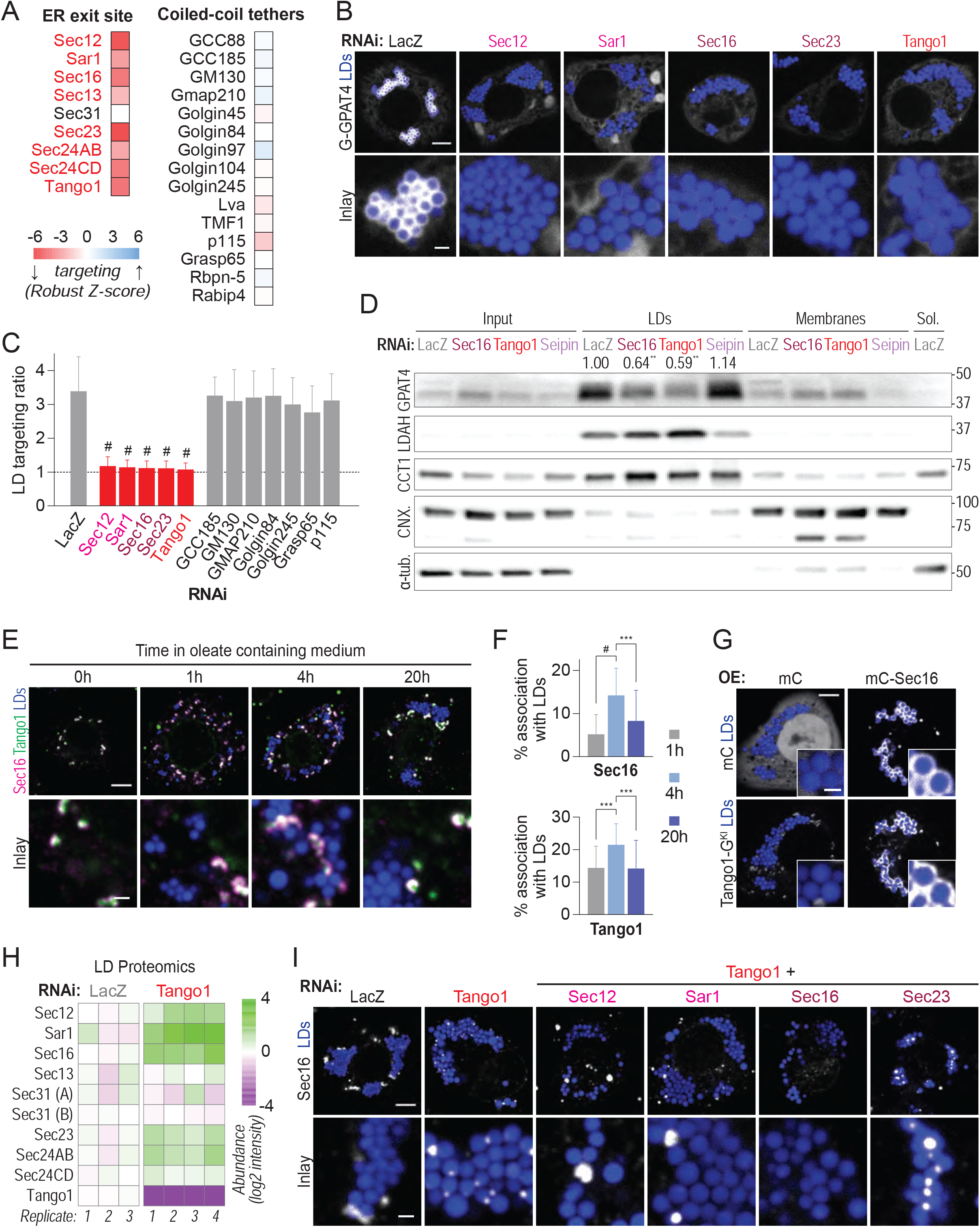
ERES organizers associate with LDs and are required for late ERTOLD targeting. (A) Heatmap of robust Z-scores for ER exit site organizers and coiled-coil tethers from the imaging screen. Genes of which knockdown results in robust Z-score < -2.5 are highlighted in red. (B) Depletion of ERES components abolishes endogenous GPAT4 targeting to LDs. Confocal imaging of live EGFP-GPAT4 endogenous knock-in cells upon RNAi of ERES components, followed by a 20-hr incubation in oleate-containing medium. LDs were stained with MDH. Representative images are shown. Scale bar, 5 and 1 μm (inlay). (C) Bar graph showing targeting ratios from the imaging experiment in (B), including select coiled-coil tethers (images not shown). Mean ± SD, n = 15–20 cells from 2–3 independent experiments. One-way ANOVA, #p<0.0001, compared to LacZ. (D) Depletion of ERES components reduces GPAT4 amount in LD fractions. Western blot analysis of fractions of wildtype *Drosophila* S2 R+ cells upon RNAi, followed by a 20-hr incubation in oleate-containing medium. Indicated on the left-hand side of the blot are protein targets of the primary antibodies, and on the right-hand side are ladder positions. Input = whole-cell lysate, sol = soluble fraction. Quantifications of GPAT4 bands in LD fractions relative to LacZ control are indicated (mean, n = 3 independent experiments). One-way ANOVA, **p<0.01, compared to LacZ. (E) ERES components Sec16 and Tango1 associate spatially with LDs 4 hours after LD induction. Immunofluorescence of Sec16 and Tango1 in wildtype cells at given timepoints after 1 mM oleic acid treatment. Scale bar, 5 and 1 μm (inlay). (F) Bar graph showing percentages of Sec16 or Tango1 puncta in association with LDs over time, calculated in 3-dimensional space per cell from the imaging experiment in (E). Mean ± SD, n = 35–40 cells from 2 independent experiments. One-way ANOVA, ***p<0.001, #p<0.0001. (G) Overexpressed Sec16 localizes around LDs and recruits endogenous Tango1 to LDs. Confocal imaging of live Tango1-EGFP endogenous knock-in cells upon transient transfection of mCherry or mCherry-Sec16 constructs, followed by a 20-hr incubation in oleate-containing medium. Scale bar, 5 and 1 μm (inlay). (H) ERES components enrich in LD fractions upon Tango1 depletion. Heatmap for abundance of ERES organizers in LD fractions upon LacZ vs. Tango1 RNAi, as measured by mass spectrometry and normalized to LacZ control. (I) Sec16 strongly co-localizes around LDs upon Tango1 depletion. Immunofluorescence of Sec16 in wildtype cells upon RNAi of Tango1 or Tango1 plus another ERES component, followed by a 20-hr incubation in oleate-containing medium. Scale bar, 5 and 1 μm (inlay).

Depletion of ERES components from cells that expressed endogenously tagged GPAT4 with additional dsRNA designs showed they are required for GPAT4 targeting to LDs, unlike coiled-coil tethering proteins (**Figures 4B and 4C**). This effect was specific to late ERTOLD targeting (Ldsdh1 and HSD17B11; **Figures S5A-D**) and did not affect LD targeting of early ERTOLD cargoes (LDAH and Ubxd8; **Figure S5E**) or CYTOLD proteins (Lsd1, CGI-58, and CCT1; **Figure S5F**). Depletion of Sec16 or Tango1 (but not of controls LacZ or seipin) also reduced abundance of GPAT4 purified with LDs in subcellular fractionation experiments, unlike LDAH or CCT1 (**Figure 4D**).

### ER exit site proteins localize to LDs

To test if ERES are involved in late ERTOLD, we analyzed the localization of Tango1 and Sec16 with respect to LD maturation by immunofluorescence. Many ERES were not localized to LDs, presumably because they operate in canonical protein export from the ER to the Golgi apparatus (**Figures 4E and 4F**). However, some ERES localized to apparent contact sites of the ER and LDs. Importantly, the association between ERES and LDs increased significantly and transiently around the time of late ERTOLD, approximately 4 hours after initiating oleic acid treatment. This suggests that ERES function in both the export of secretory proteins to the Golgi apparatus and late ERTOLD targeting to LDs. In agreement with a direct role of ERES at LDs, overexpressed Sec16 (fused with mCherry) localized around LDs and recruited endogenous Tango1 (fused with EGFP) to LDs (**Figure 4G**), indicating that Sec16 may act upstream of Tango1 at LDs. Additionally, expression of a tagged dominant-negative version of Sar1 (the GDP-restricted T34N mutant) formed ring-like signals around LDs and impaired targeting of endogenously tagged GPAT4 to LDs (**Figures S2B and S2C**).

The transiently increased association of LDs and Sec16 and Tango1 was accompanied by an increase in the number of ERES per cell at 4 hours after initiating oleic acid treatment (**Figures 4E-F and S5G**). This suggests a role for ERES at LDs, independent of the canonical secretory pathway. To better understand how the different ERES components organize around LDs, we depleted cells of the key component Tango1 and tested the association of other ERES components with LDs by mass spectrometry. Strikingly, abundance of ERES components Sec12, Sar1, Sec16, and Sec23 was robustly increased in LD fractions from cells depleted of Tango1 compared with LacZ control (**Figure 4H**). Consistent with this, immunofluorescence microscopy showed increases in Sec16 puncta association with LDs upon Tango1 RNAi (**Figure 4I**, two left-most panels). Similarly, Tango1 depletion increased association of the overexpressed constitutively active Sar1 H74G mutant (defective in GTP hydrolysis) and Sec23 with LDs by confocal microscopy (**Figure S5H**). Thus, depletion of the ERES protein Tango1 appears to increase Sar1, Sec23, and Sec16 localization to ER-LD contact sites.

To assess if Sec16 recruitment to LDs requires other ERES factors, we depleted Sec12, Sar1, or Sec23 on top of Tango1 and tested Sec16 association with LDs. Sec23 depletion did not affect Tango1 depletion-mediated Sec16 recruitment to LDs, but Sec12 or Sar1 depletion (with Tango1 depletion) abolished Sec16 recruitment to LDs (**Figure 4I**). Similarly, localization of overexpressed, tagged Sec16 around LDs required Sec12 and Sar1 but not Sec23 (**Figure S5I**). This contrasts with the previous finding that Sec16 localization to canonical ERES was independent of Sar1 (Ivan et al., 2008) and suggests differences in the ERES organization for ERTOLD and Golgi protein targeting.

### Seipin restricts late ERTOLD cargoes from accessing LDs during their formation

Late ERTOLD targeting results in an apparent paradox: why do late ERTOLD cargo proteins stay in the ER not binding to LDs when they form? At LD formation sites, seipin forms a large complex with 20–24 transmembrane domains (depending on species) forming a 10–15-nm ring that appears to localize around the budding neck of forming LDs (Arlt et al., 2021; Sui et al., 2018; Yan et al., 2018). This structure suggests that the seipin complex may restrict ER proteins from accessing forming LDs. To test this possibility, we compared targeting kinetics of early and late ERTOLD proteins in cells lacking seipin. Unlike in wildtype cells (**Figure 1**), all analyzed early (LDAH, Ubxd8) and late ERTOLD proteins (GPAT4, Ldsdh1, HSD17B11) targeted LDs as early as 30 minutes after LD induction in seipin knock-out cells (**Figure 5A**).

**Figure 5.**
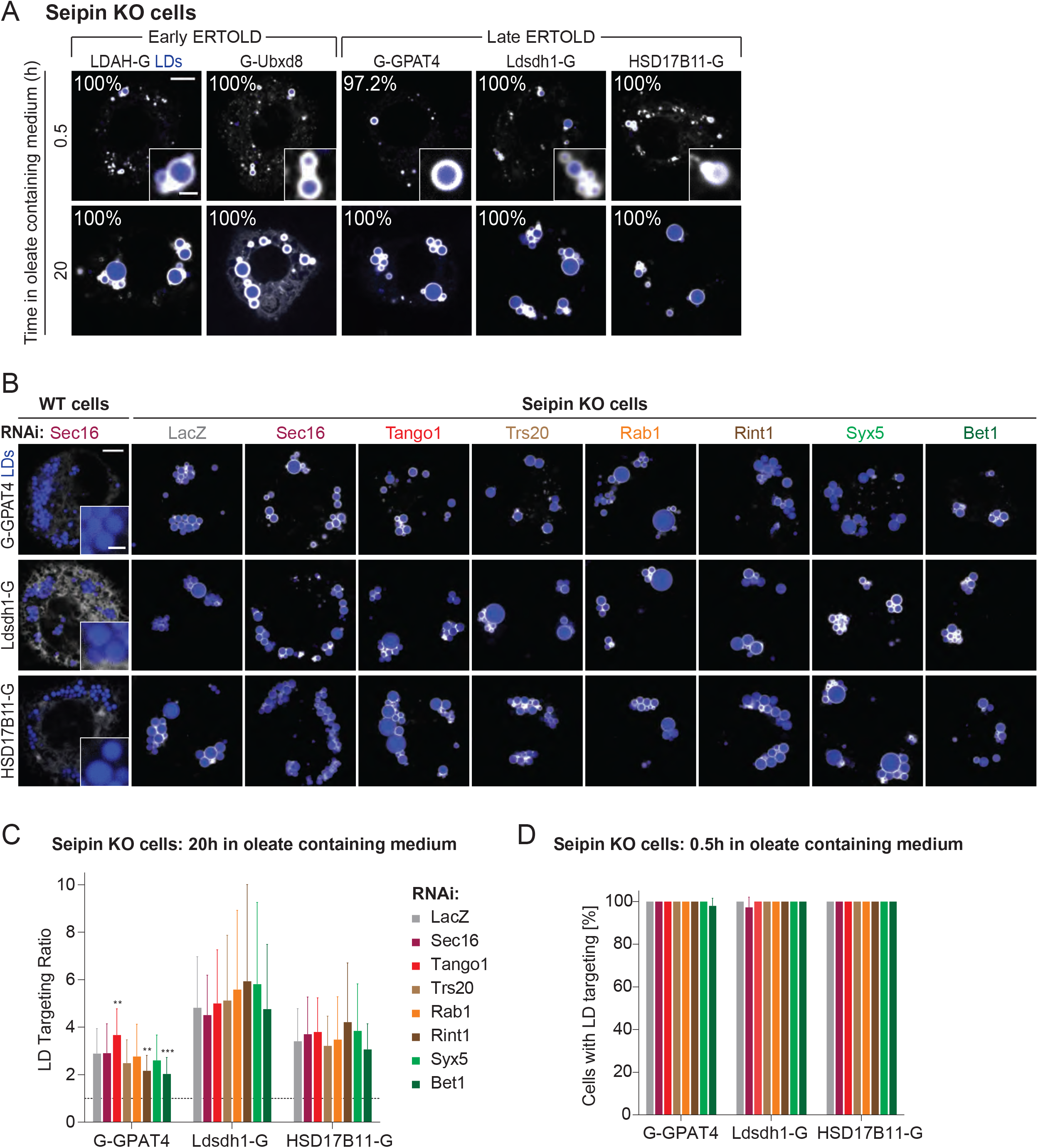
Seipin depletion allows for late ERTOLD targeting in the absence of fusion machinery or ERES components. (A) Late ERTOLD proteins target LDs early in absence of seipin. Confocal imaging of live Seipin knockout cells transiently transfected with EGFP-tagged constructs at given timepoints after 1 mM oleic acid treatment. LDs were stained with MDH. Representative images are shown. Percentage of cells with LD targeting are indicated (mean, n = 3 independent experiments, 8–13 cells each). Scale bar, 5 and 1 μm (inlay). (B) Absence of seipin provides an alternative pathway for LD targeting of late ERTOLD proteins. Confocal imaging of live wildtype or Seipin knock-out cells upon RNAi of ERES or fusion-machinery components, followed by transient transfection with EGFP-tagged constructs and a 20-hr incubation in oleate-containing medium. Scale bar, 5 and 1 μm (inlay). (C) Bar graph showing targeting ratios from the imaging experiment in (B). Mean ± SD, n = 35– 50 cells from 3 independent experiments. One-way ANOVA, **p<0.01, ***p<0.001, compared to LacZ. (D) Bar graph showing percentages of cells with LD targeting after a 0.5-hr incubation in oleate-containing medium. Representative images are shown in **Figure S6A**.

When we depleted one of the ERES or ERTOLD membrane fusion machinery proteins (Sec16, Tango1, Trs20, Rab1, Rint1, Syx5 or Bet1) in seipin knock-out cells, we found that impaired localization of late ERTOLD targeting cargoes was rescued, compared with the defect found in wildtype cells depleted for these factors (**Figures 3A-B, 4B-C, and 5B-C**). Specifically, late ERTOLD proteins targeted LDs during their formation in the absence of seipin, even when the ERES or ERTOLD membrane fusion machinery proteins were depleted (**Figures 5D and S6A**). Importantly, their depletion did not alter the formation of endogenous seipin foci near LDs (**Figure S6B**). This shows that seipin functions as a negative regulator of protein targeting to forming LDs, restricting the access of specific ERTOLD cargoes. This also indicates that depletion of late ERTOLD machinery does not impair the ability of late ERTOLD cargoes to move to LDs but instead abolishes their path to LDs, and the absence of seipin provides an alternative route for late ERTOLD targeting.

### Systematic identification of late ERTOLD cargoes

Identification of the machinery for late ERTOLD protein targeting enabled us to systematically identify proteins that depend on this trafficking route to access LDs. For this, we individually depleted members of the ERES or ERTOLD membrane fusion machinery (Tango1, Trs20, Rab1, Rint1, Syx5 or Bet1) from cells, isolated LDs, and analyzed their composition by mass spectrometry-based proteomics. Seipin knock-out cells and RNAi treatment targeting LacZ were included as controls. As expected, protein levels of each factor targeted by RNAi were dramatically reduced (**Figure S7A**). Filtering the data for the set of LD proteins (Krahmer et al., 2013) with two or four consecutive predicted transmembrane domains that may form membrane-embedded hairpins, we found ∼10 proteins whose abundance in LD fractions was reduced by depletion of late ERTOLD targeting machinery components. These proteins included LPCAT, ACSL5, ReepA, and DHRS7B, in addition to GPAT4 (**Figure 6A**).

**Figure 6.**
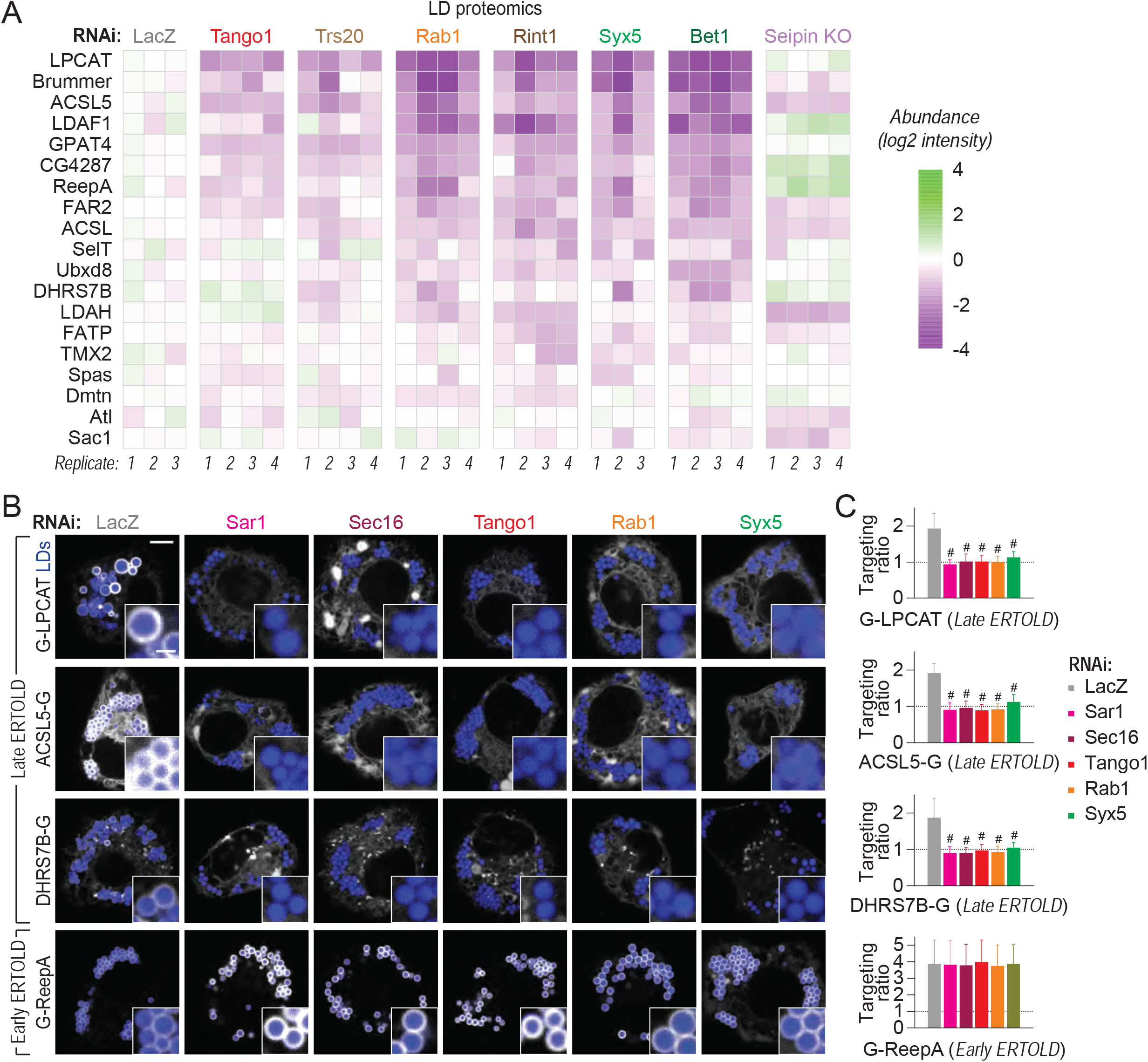
LD proteomics reveal additional late ERTOLD cargoes. (A) Heatmap of abundance of potential ERTOLD proteins in LD fractions upon depletion of the late ERTOLD machinery components or seipin (compared to LacZ control), as measured by mass spectrometry. (B) LPCAT, ACSL5, and DHRS7B require ERES or fusion-machinery components for LD targeting. Confocal imaging of live wildtype cells upon RNAi of ERES or fusion-machinery components, followed by transient transfection with EGFP-tagged constructs and a 20-hr incubation in oleate-containing medium. LDs were stained with MDH. Scale bar, 5 and 1 μm (inlay). (C) Bar graph showing targeting ratios from the imaging experiment in (B). Mean ± SD, n = 30- 50 cells from 3 independent experiments. One-way ANOVA, #p<0.0001, compared to LacZ.

To confirm their targeting and dependence on late ERTOLD membrane fusion machinery, we expressed fluorescently tagged versions of these proteins in cells lacking Sar1, Sec16, Tango1, Rab1, or Syx5 and compared them with LacZ controls. As with the proteomics results, we found that LD localization of ACSL5, DHRS7B, and LPCAT was strongly impaired by depletion of the late ERTOLD machinery (**Figures 6B and 6C**). ReepA targeting was not impaired, but it targeted LDs early instead (**Figure S7 and S7C**).

## DISCUSSION

Here we addressed a major gap in our understanding of protein targeting in eukaryotic cells: how do proteins localize from the bilayer membrane of the ER to the monolayer of LDs? We delineate two distinct mechanisms for ERTOLD targeting: early ERTOLD targeting with proteins transiting through LDACs during LD formation, and late ERTOLD targeting via independently established membrane bridges that connect the ER with mature LDs. We identified the membrane fusion machinery for late ERTOLD targeting and a requirement for ERES machinery in mediating late ERTOLD targeting. We uncovered a spatial association of LDs with many of these machinery components, including ERES. Additionally, we found numerous cargoes of early and late ERTOLD pathways.

Previously, we and others showed that some artificial ER-embedded hairpins, such as *LiveDrop* (Wang et al., 2016) or HPos (Kassan et al., 2013), access LDs during their formation. The LDAC component LDAF1, which appears to contain two hairpins, also accesses LDs during formation (Chung et al., 2019). We now find that early ERTOLD targeting occurs for other cellular proteins, including Ubxd8, LDAH, and ReepA. How some ER proteins, but not others, access nascent LDs during their formation is unclear. Current data suggest that the ring of seipin at LDACs restricts some but not all ERTOLD proteins from accessing nascent LDs [**Figure 7**, left-most panel; (Arlt et al., 2021; Sui et al., 2018; Yan et al., 2018)].

**Figure 7.**
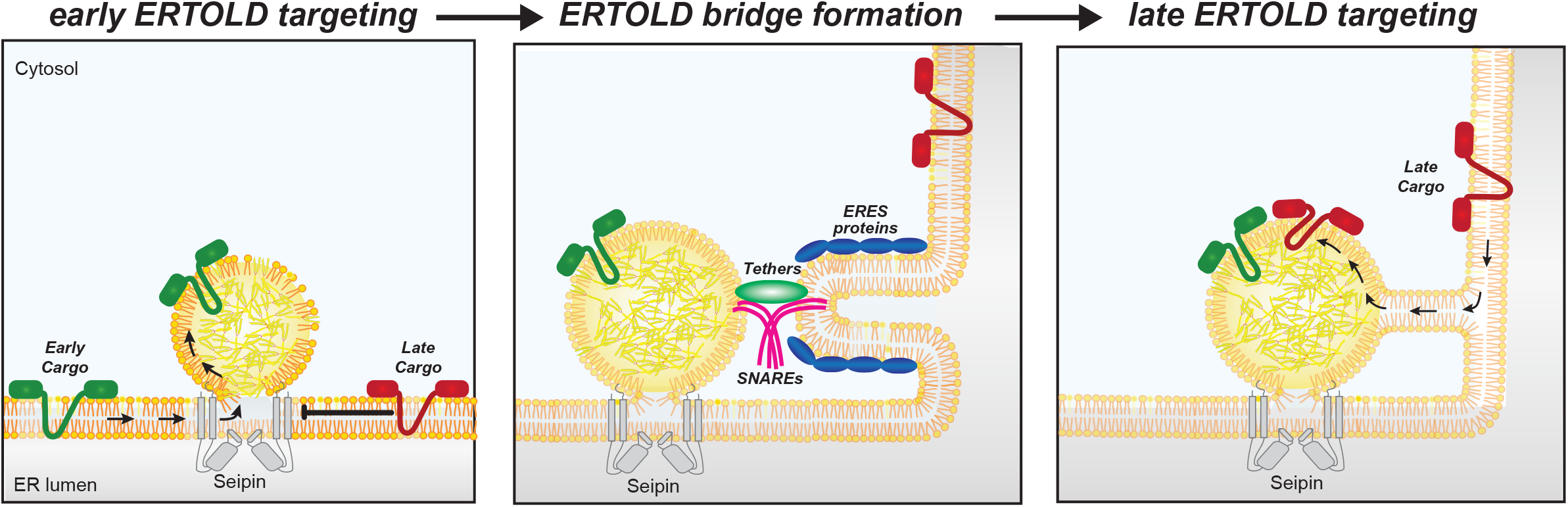
Model of ERTOLD targeting. Early cargoes can access forming LDs from the ER through the LDACs, whereas late cargoes cannot. A process mediated by the membrane-fusion machinery, including a Rab protein, membrane tethers and SNAREs, at ERES forms an ERTOLD bridge, which then allows LD targeting of late cargoes, such as GPAT4 and LPCAT, that are crucial for lipid metabolism on LD surfaces.

The barrier between the ER and forming LDs necessitates an alternative pathway for protein targeting to LDs. Studies in *Drosophila* cells suggested that GPAT4 is excluded from nascent LDs and traffics to LDs via membrane bridges (Wilfling et al., 2013). The estimated speed of GPAT4 targeting to LDs was consistent with diffusion within membranes but was too fast for vesicular trafficking (Wilfling et al., 2014). These findings suggested that cells possess machinery to generate ER-LD membrane bridges, but how the bridges are established was unknown.

By performing an unbiased screen for factors enabling late ERTOLD targeting of GPAT4, we identified components required for each step of membrane fusion, including Rab1, Rint1, Trs20 and specific SNAREs. Rab GTPases act as switches for regulating various steps of intracellular membrane trafficking (Stenmark, 2009). Specifically, Rab1 has been implicated in tethering COPII-coated vesicles from the ER to *cis*-Golgi by interacting with coil-coiled tether, such as p115 and GM130 (Allan et al., 2000; Beard et al., 2005; Moyer et al., 2001). Given the localization of Rab1 on LDs here and by others (Bersuker et al., 2018; Krahmer et al., 2013), we suspect that Rab1 acts as a molecular switch priming the LD membranes for heterotypic fusion with the ER. Alternatively, it may facilitate the extrusion of late ERTOLD cargoes at ERES (Westrate et al., 2020). Whereas p115 and GM130 were dispensable for late ERTOLD targeting, we found another membrane tethering complex component Rint1, which was proposed to help in establishing ER-LD contacts (Xu et al., 2018), to localize around LDs and be required for late ERTOLD targeting.

Trs20, which is a shared component of different TRAPP complexes (Barrowman et al., 2010), was required for late ERTOLD targeting. TRAPP complexes may act as a GEF or a tethering complex for Rab1 (Kim et al., 2016). In agreement with a role of Trs20 in activating Rab1 in late ERTOLD targeting, expression of a Rab1 dominant-negative mutant (which sequesters TRAPP complexes) abolished GPAT4 targeting to LDs (**Figure S2B and S2C**). Intriguingly, no other component of TRAPP complexes was required for late ERTOLD targeting, which argues for a specific role of Trs20 in this context. Trs20 for instance may be critical for shaping the ER adjacent to LDs for protein export, as was suggested for its human ortholog Sedlin at ERES during the export of large cargoes, such as procollagen (Brandizzi and Barlowe, 2013; Venditti et al., 2012).

The SNARES we identified as necessary for late ERTOLD targeting constitute components of a putative SNAREpin: Syx5 (Qa SNARE), membrin (Qb), Bet1 (Qc) and Ykt (R). The capacity of this combination to fuse the ER membrane with a LD has not yet been tested, and which of the SNAREs may act on the side of LDs is unclear. Ykt6 is a good candidate as it is anchored to the membrane via a lipid modification (Fukasawa et al., 2004), rather than a transmembrane domain which would be incompatible with localizing to the LD monolayer. Also, Ykt6 was identified as a potential LD protein in systematic studies of LD composition (Bersuker et al., 2018; Krahmer et al., 2018).

Unexpectedly, our screen also identified many ERES components, including Sec12, Sar1, Sec16, and Tango1 (Budnik and Stephens, 2009; Liu et al., 2017; Raote et al., 2018), as required for late ERTOLD protein targeting. Consistent with previous observations that ERES localize near LDs (Soni et al., 2009), we found transient association between ERES and LDs around the time late ERTOLD targeting begins. Thus, an ERES analogous domain may function in establishing late ERTOLD targeting bridges. This might explain why ATGL colocalizes with ERES markers when the COPI machinery is impaired (Soni et al., 2009). ERES domains functioning in secretory pathway or late ERTOLD targeting may compete for some of the same components since the number of Sec16 and Tango1 puncta increases in response to LD induction and many ERES components re-localize to sites around LDs when the ERES organizing protein Tango1 is depleted.

Arf1/COPI coatomer proteins are required for LD targeting of GPAT4 and ATGL (Beller et al., 2008; Soni et al., 2009; Wilfling et al., 2014). A current model for the function of Arf1/COPI proteins posits that they localize directly on LDs, modulating their surface properties (Thiam et al., 2013b; Wilfling et al., 2014). Although we do not know yet if they act before or after SNAREs, these proteins may be needed to modify the LD surface to accommodate fusion factors, such as Rab1 or Ykt6.

An attractive unifying model based on these and other findings (Raote et al., 2017, 2018; Weigel et al., 2021) is that the ER forms tubular carriers at ERES that connect to different target organelles. In the case of secretory trafficking, formation of such tubes or tunnels allows for secretion of large cargoes, such as collagens or lipoproteins (Raote et al., 2018; Santos et al., 2016), to the Golgi apparatus. In our model of late ERTOLD, tubular structures formed at ERES could instead fuse with LDs to form bridges between both organelles (**Figure 7**, middle panel). The result of the fusion reaction is that the cytosolic leaflet of the ER bilayer becomes contiguous with the LD monolayer surface, allowing membrane-embedded proteins to traverse between the organelles (**Figure 7**, right-most panel). Late ERTOLD proteins may then accumulate on LDs because this location is energetically favorable over the ER (Olarte et al., 2020).

### Limitations of the Study

We previously reported microscopy evidence showing membrane bridges between the ER and LDs mediating protein targeting (Wilfling et al., 2013, 2014). Although evidence suggests that the identified membrane fusion and ERES proteins act directly at LDs to establish such ER-LD membrane bridges, direct visualization of this reaction is not available. Likely, the fusion reactions are fast, and only a few events are needed to mediate targeting. Moreover, the machinery used in this reaction is shared with other membrane trafficking reactions. Thus, like other membrane trafficking events, the establishment of membrane bridges is difficult to observe, and future work will have to focus on the biochemical dissection of the reaction, as well as visualization of membrane bridges.

The pathway of late ERTOLD targeting appear to be evolutionarily conserved between flies and humans, and some of the hits of our screen, such as Arf1/COPI, are required for ERTOLD targeting in both systems (Beller et al., 2008; Soni et al., 2009; Wilfling et al., 2014). It seems likely that late ERTOLD targeting operates in mammalian cells, inasmuch as salient features of cell biology are mostly conserved between organisms. This, however, will require further testing. Importantly, proteins involved in metabolic diseases, such as HSD17B13 (an orthologue of late ERTOLD cargo HSD17B11 in this study), may utilize this pathway, highlighting the importance of understanding the mechanisms of ERTOLD targeting for possible therapeutics.

## ACKNOWLEDGMENTS

We thank the members of Farese & Walther laboratory, Drs. Timothy Mitchison (HMS), Norbert Perrimon (HMS, HHMI), and Adrian Salic (HMS) for helpful discussions. We also thank Drs. Mathias Beller (Heinrich Heine Universitat Dusseldorf, Germany), Sally Horne-Badovinac (University of Chicago), Luke Lavis (HHMI), Catherine Rabouille (Hubrecht Institute, Netherlands), and Norbert Perrimon for reagents; Rong Tao, Dr. Jonathan Zirin, Dr. Yanhui Hu, Dr. Donghui Yang-Zhou, Shannon Knight, and Gabriel Amador (DRSC, Department of Genetics, HMS) for technical support with screening; the Nikon Imaging Center (HMS) for imaging support; the Flow Cytometry Core (Department of Immunology, HMS) for the support with cell sorting; and Garry Howard for editorial assistance.

This work was supported by the grants from the National Institutes of Health R01GM097194 (T.C.W), 5TL1TR001101 (J.S.), and T32GM007753 (J.S.). J.S. received support from the American Heart Association Predoctoral Fellowship and the Aramont Fund for Emerging Science Research. T.C.W. is a Howard Hughes Medical Institute investigator. A.M. is a Helen Hay Whitney Foundation Postdoctoral Fellow. DRSC is supported by NIGMS P41 GM132087.

## AUTHOR CONTRIBUTIONS

J.S., R.V.F., and T.C.W. conceived the project. J.S., R.V.F., and T.C.W. designed the screen, and S.M. and M.C. provided additional input. M.C. and J.S. built the pipeline for screen image analysis. J.S. performed and analyzed the screen. J.S. performed and analyzed most of the experiments. A.M. created Tango1 knock-in cell line. C-W.L. performed co-expression of LDAH and GPAT4 in cells. Z.W.L. performed mass spectrometry analyses. C-H.L. performed spatial analysis of LDs and ERES. J.S., R.V.F., and T.C.W. wrote the manuscript. All authors discussed the results and contributed to the manuscript.

## DECLARATION of INTRESTS

The authors declare no competing interests.

## SUPPLEMENTAL FIGURE LEGENDS

**Figure S1.**
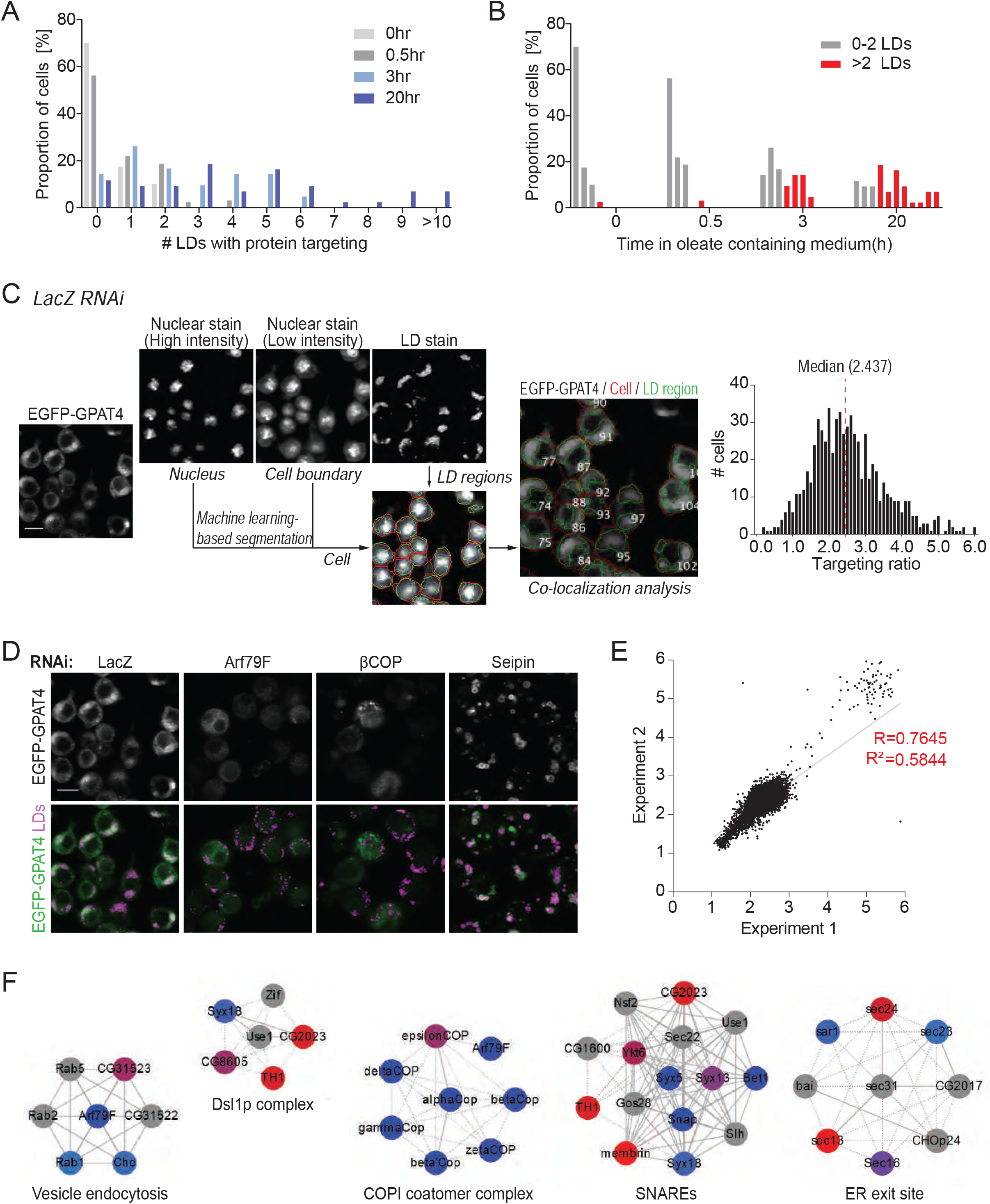
HSD17B11 targeting to LDs and an imaging screen to identify factors required for GPAT4 targeting to LDs, related to Figures 1 and 2. (A) Bar graph showing percentage of cells for a given number of LDs with HSD17B11-EGFP targeting over time after 1 mM oleic acid treatment. Representative images are shown in Figure 1B. (B) Representation of (A) with respect to time in oleate-containing medium. (C) Schematic diagram for the calculation of targeting ratios in the imaging screen. Sample images from a well containing LacZ control are shown. Machine learning–based segmentation methods were used to segment individual cells and regions of LDs from the nuclear and LD stains, which are superimposed on EGFP-GPAT4 channel to calculate LD targeting ratio for each cell. Median of the targeting ratios from all segmented cells in eight different fields of the well is reported as the final readout. (D) Representative images for screen controls. LacZ RNAi has no effect on GPAT4 targeting to LDs. Depletion of Arf79F or βCOP reduces GPAT4 targeting to LDs. Depletion of seipin increases GPAT4 targeting to LDs. Scale bar, 10 μm. (E) Genome-scale screen results are reproducible. Scatter plot showing targeting ratios from 2 independent genome-scale screen experiments. Linear regression line is shown in grey. R and R2 values are indicated. (F) Protein complexes enriched among hits required for GPAT4 targeting to LDs (defined as robust Z-score < -2.5) using COMPLEAT (Vinayagam et al., 2013). Node color: Blue to red (lowest to highest robust Z-scores), grey (non-hits, robust Z-score > 2.5). Line type: solid (known interaction), dashed (known interaction among orthologs in another species). p-value < 0.01 for all complexes shown.

**Figure S2.**
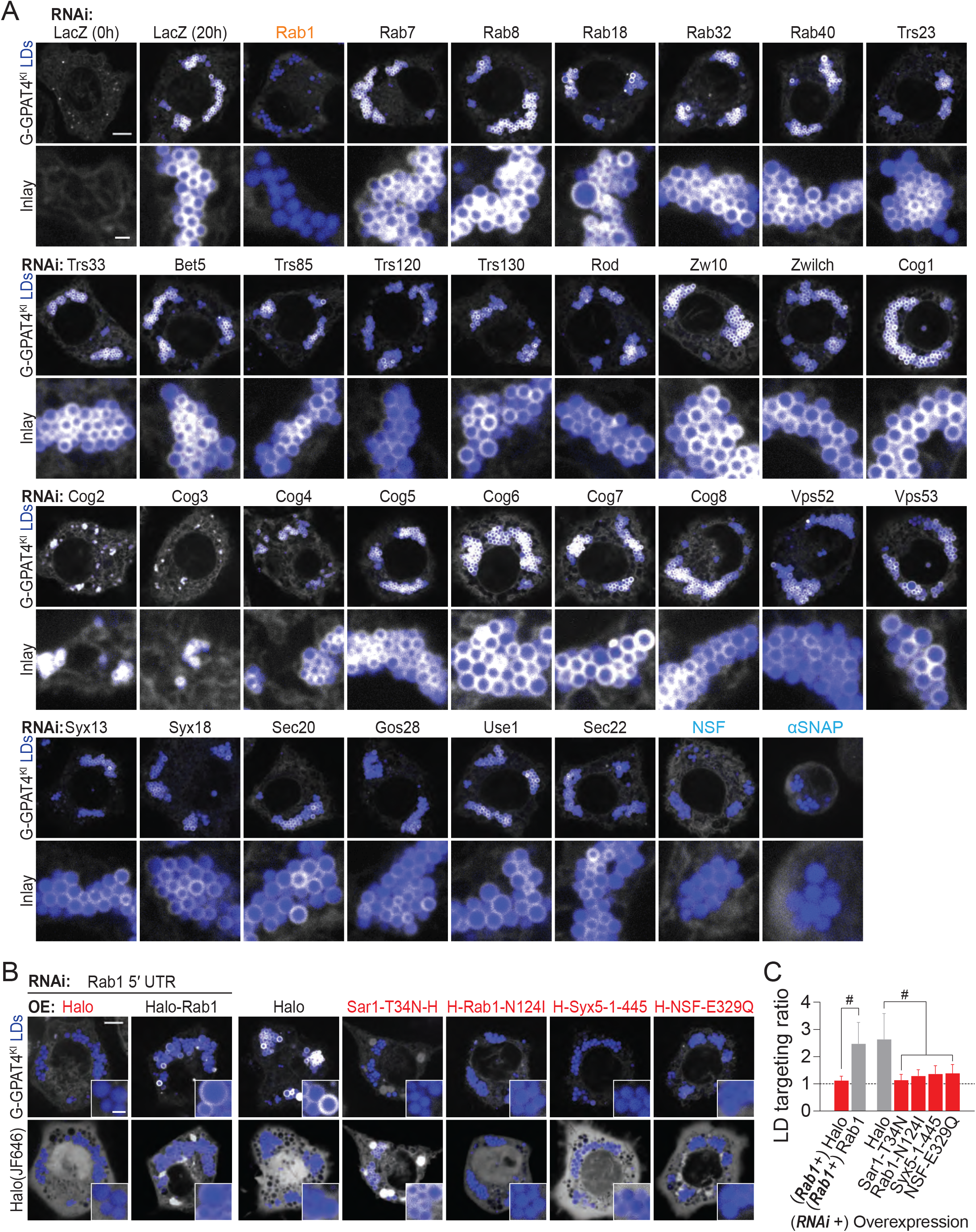
Specific membrane-fusion regulator, tether, and SNAREs are required for GPAT4 targeting to LDs, related to Figure 3. (A) Depletion of specific Rab, tethering-complex components, and SNAREs abolishes LD targeting to LDs. Confocal imaging of live EGFP-GPAT4 endogenous knock-in cells upon RNAi of membrane fusion machinery components, 20 h after 1-mM oleic acid treatment (except for 0 h timepoint for LacZ RNAi). LDs were stained with MDH. Representative images are shown. Scale bar, 5 and 1 μm (inlay). Quantification of targeting ratios is shown in Figure 3B. (B) Effect of Rab1 depletion on LD targeting of GPAT4 is rescued by wildtype Rab1 expression, and the expression of dominant-negative Sar1, Rab1, Syx5, and NSF mutants impairs GPAT4 targeting to LDs. Confocal imaging of live EGFP-GPAT4 endogenous knock-in cells upon RNAi and/or overexpression of Halo tagged constructs. H = Halo tag. Scale bar, 5 and 1 μm (inlay). (C) Bar graph showing targeting ratios from the imaging experiment in (B). Mean ± SD, n = 40– 60 cells from 3 independent experiments. One-way ANOVA, #p<0.0001.

**Figure S3.**
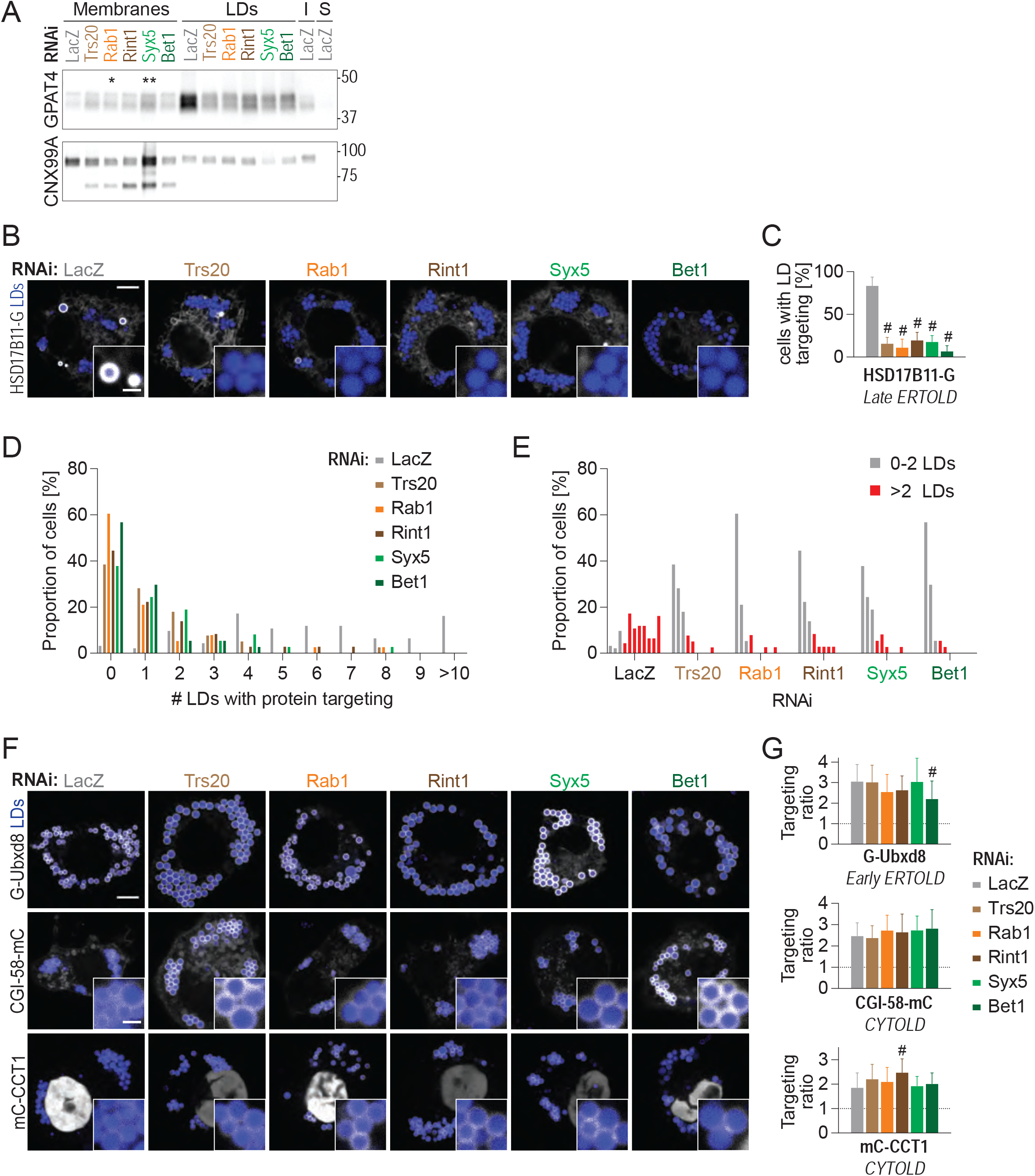
Membrane-fusion regulator, tether, and SNAREs are required specifically for late ERTOLD targeting, related to Figure 3. (A). Western blot analysis of fractions of wildtype cells upon RNAi of fusion machinery components, followed by a 20-hr incubation in oleate-containing medium. GPAT4 amount increases in membrane fractions but decrease in LD fractions upon depletion of membrane-fusion machinery components. Indicated on the left-hand side of the blot are protein targets of the primary antibodies, and on the right-hand side are ladder positions. I or Input = whole-cell lysate, S = soluble fraction. Intensities of GPAT4 bands in membrane fractions relative to LacZ control (1.00) were: Trs20 (2.25 ± 0.58), Rab1* (2.62 ± 1.04), Rint1 (2.11 ± 0.25), Syx5** (3.07 ± 0.40), and Bet1 (2.17 ± 0.28) (mean ± SD, n = 3 independent experiments). One-way ANOVA, *p<0.05, **p<0.01, compared to LacZ. (B) HSD17B11 requires membrane-fusion machinery components for targeting from the ER to LDs. Confocal imaging of live wildtype cells transiently transfected with HSD17B11-EGFP upon RNAi of fusion machinery, followed by a 20-hr incubation in oleate-containing medium. Scale bar, 5 and 1 μm (inlay). (C) Bar graph showing percentage of cells with LD targeting (defined as those with >2 LDs with protein targeting) from the imaging experiment in (B). Mean ± SD, n = 3 independent experiments (12–17 cells each). One-way ANOVA, #p<0.0001, compared to LacZ. (D) Bar graph showing percentage of cells for a given number of LDs with HSD17B11-EGFP targeting upon RNAi of membrane-fusion machinery. Representative images are shown in (B). (E) Representation of (D) with respect different RNAi. (F) Depletion of specific Rab, membrane-tethering complex components, and SNAREs does not impair targeting of Ubxd8 (early ERTOLD) or CGI-58 and CCT1 (CYTOLD). Confocal imaging of live wildtype cells transiently transfected with EGFP- or mCherry-tagged constructs upon RNAi of fusion machinery, followed by a 20-hr incubation in oleate-containing medium. Scale bar, 5 and 1 μm (inlay). (G) Bar graph showing targeting ratios from the imaging experiment in (G). Mean ± SD, n = 30– 60 cells from 3 independent experiments. For mCherry-CCT1, nuclear signal was excluded from the calculation. One-way ANOVA, #p<0.0001, compared to LacZ.

**Figure S4.**
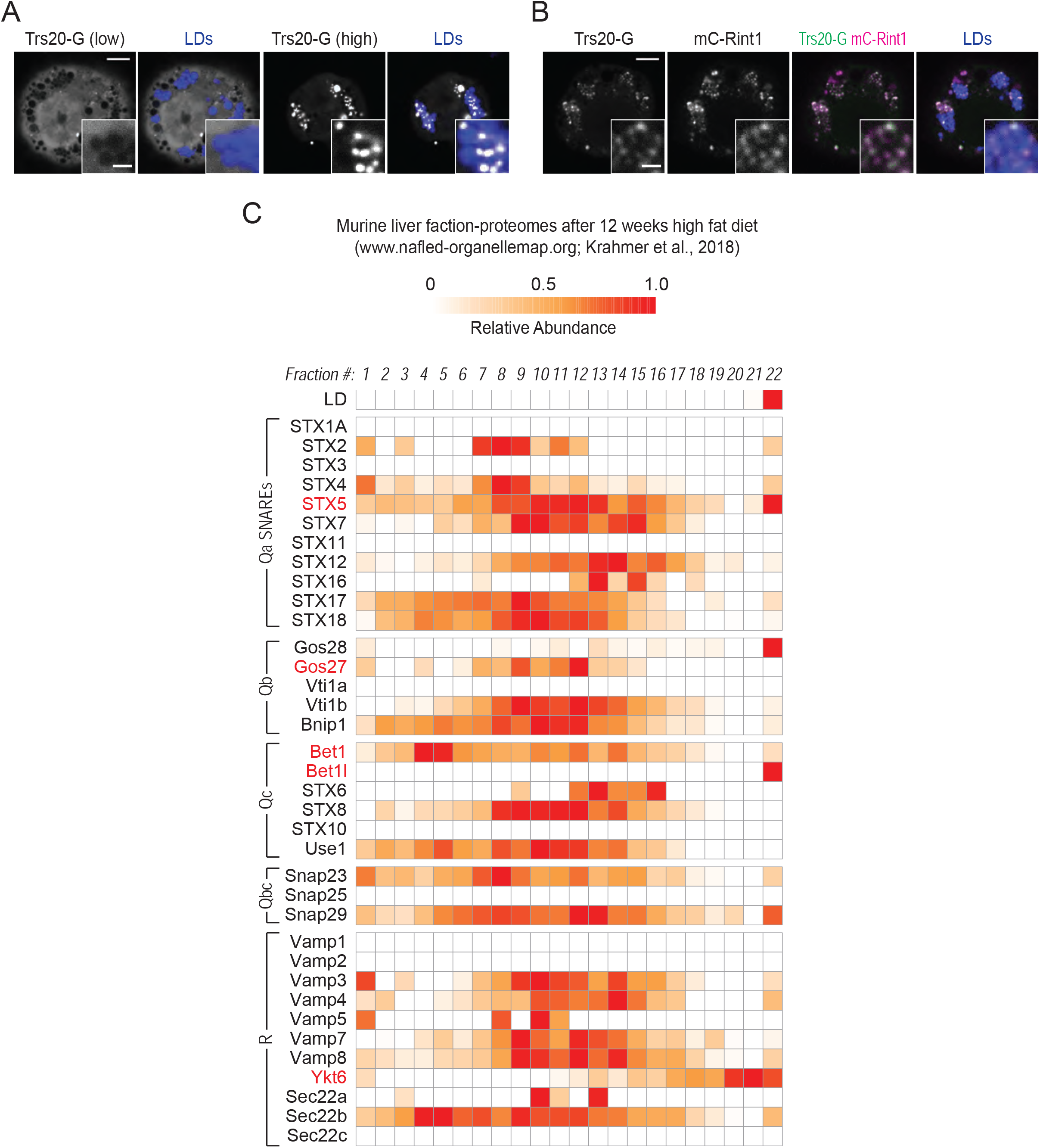
Trs20 and specific SNAREs associate with LDs, related to Figure 3. (A) Trs20 forms punctate localizations around LDs when highly overexpressed. Localization of transiently transfected Trs20-EGFP with respect to LDs. “Low” and “high” indicate the degree of overexpression, based on fluorescence intensity. Scale bar, 5 and 1 μm (inlay). (B) Overexpressed Rint1 recruits Trs20 to LDs. Localization of transiently co-transfected Trs20-EGFP and mCherry-Rint1 with respect to LDs. Scale bar, 5 and 1 μm (inlay). (C) Specific SNAREs are enriched in the LD fraction of murine fatty liver [data mined from nafld-organellemap.org; (Krahmer et al., 2018)]. In this study, mice were subjected to 12 weeks of high-fat diet, their livers were harvested and separated into 22 fractions using a sucrose gradient, and proteomes of the fractions were analyzed with mass spectrometry. First row shows the organellar migration pattern for LDs based on protein correlation profiling. SNAREs are classified according to their classes, and the predicted orthologs of SNAREs required for late ERTOLD targeting (Syx5, membrin, Bet1, and Ykt6) are highlighted in red.

**Figure S5.**
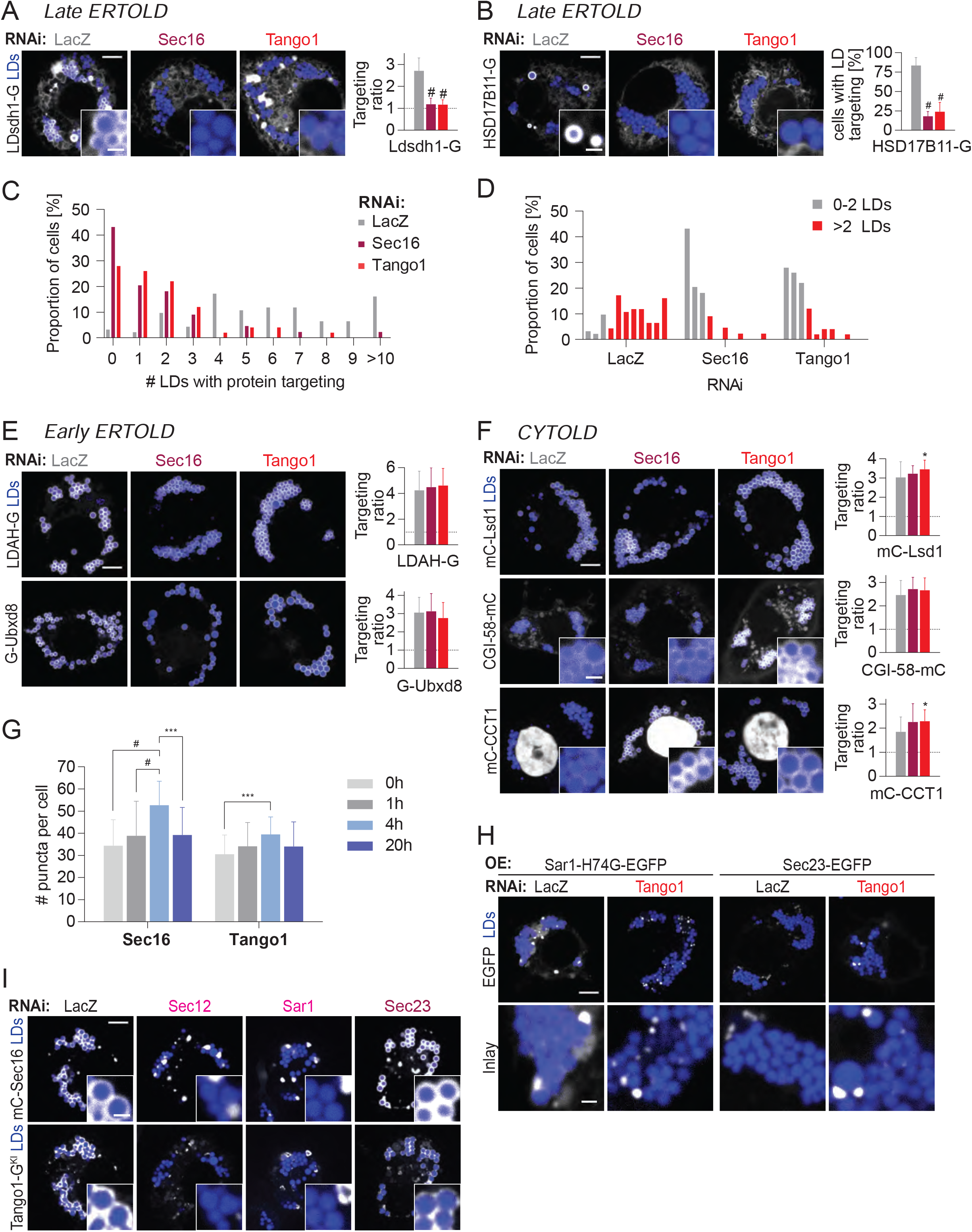
ERES increase in number upon LD induction and are specifically required for late ERTOLD targeting, related to Figure 4. (A) Sec16 and Tango1 are required for LD targeting of Ldsdh1. Confocal imaging of live wildtype cells upon RNAi of ERES components, followed by transient transfection with Ldsdh1-EGFP and a 20-hr incubation in oleate-containing medium. Scale bar, 5 and 1 μm (inlay). Bar graph shows targeting ratios calculated from the images. Mean ± SD, n = 40–50 cells from 3 independent experiments. One-way ANOVA, #p<0.0001, compared to LacZ. (B) Sec16 and Tango1 are required for LD targeting of HSD17B11. Confocal imaging of live wildtype cells upon RNAi of ERES components, followed by transient transfection with HSD17B11-EGFP and a 20-hr incubation in oleate-containing medium. Scale bar, 5 and 1 μm (inlay). Bar graph shows percentage of cells with LD targeting (defined as those with >2 LDs with protein targeting). Mean ± SD, n = 3 independent experiments (14–17 cells each). One-way ANOVA, #p<0.0001, compared to LacZ. (C) Bar graph showing percentage of cells for a given number of LDs with HSD17B11-EGFP targeting upon RNAi of ERES components. Representative images are shown in (B). (D) Representation of (C) with respect different RNAis. (E–F) Sec16 and Tango1 are dispensable for LD targeting of LDAH and Ubxd8 (early ERTOLD) and of Lsd1, CGI-58, and CCT1 (CYTOLD). Confocal imaging of live wildtype cells upon RNAi of fusion machinery, followed by transient transfection of EGFP or mCherry tagged constructs and a 20-hr incubation in oleate-containing medium. Scale bar, 5 and 1 μm (inlay). Bar graph shows targeting ratios calculated from the images. For mCherry-CCT1, nuclear signal was excluded from the calculation. Mean ± SD, n = 30–50 cells from 3 independent experiments. One-way ANOVA, *p<0.05, compared to LacZ. (G) LD induction transiently increases the number of Sec16 and Tango1 puncta. Bar graph showing the number of Sec16 or Tango1 puncta in 3-dimensional space per cell over time in oleate-containing medium. Representative images are shown in Figure 4E. Mean ± SD, n = 35–40 cells from 2 independent experiments. One-way ANOVA, ***p<0.001, #p<0.0001. (H) Overexpressed Sar1-H74G and Sec23 accumulate around LDs upon Tango1 depletion. Confocal imaging of live wildtype cells upon RNAi of LacZ or Tango1, followed by transient transfection of EGFP tagged ERES components and a 20-hr incubation in oleate-containing medium. Scale bar, 5 and 1 μm (inlay). (I) Localization of Sec16 around LDs and its recruitment of endogenous Tango1 require Sec12 and Sar1 but not Sec23. Confocal imaging of live Tango1-EGFP endogenous knock-in cells upon RNAi of ERES components followed by transient transfection with an mCherry-Sec16 construct and a 20-hr incubation in oleate containing medium. Scale bar, 5 and 1 μm (inlay).

**Figure S6.**
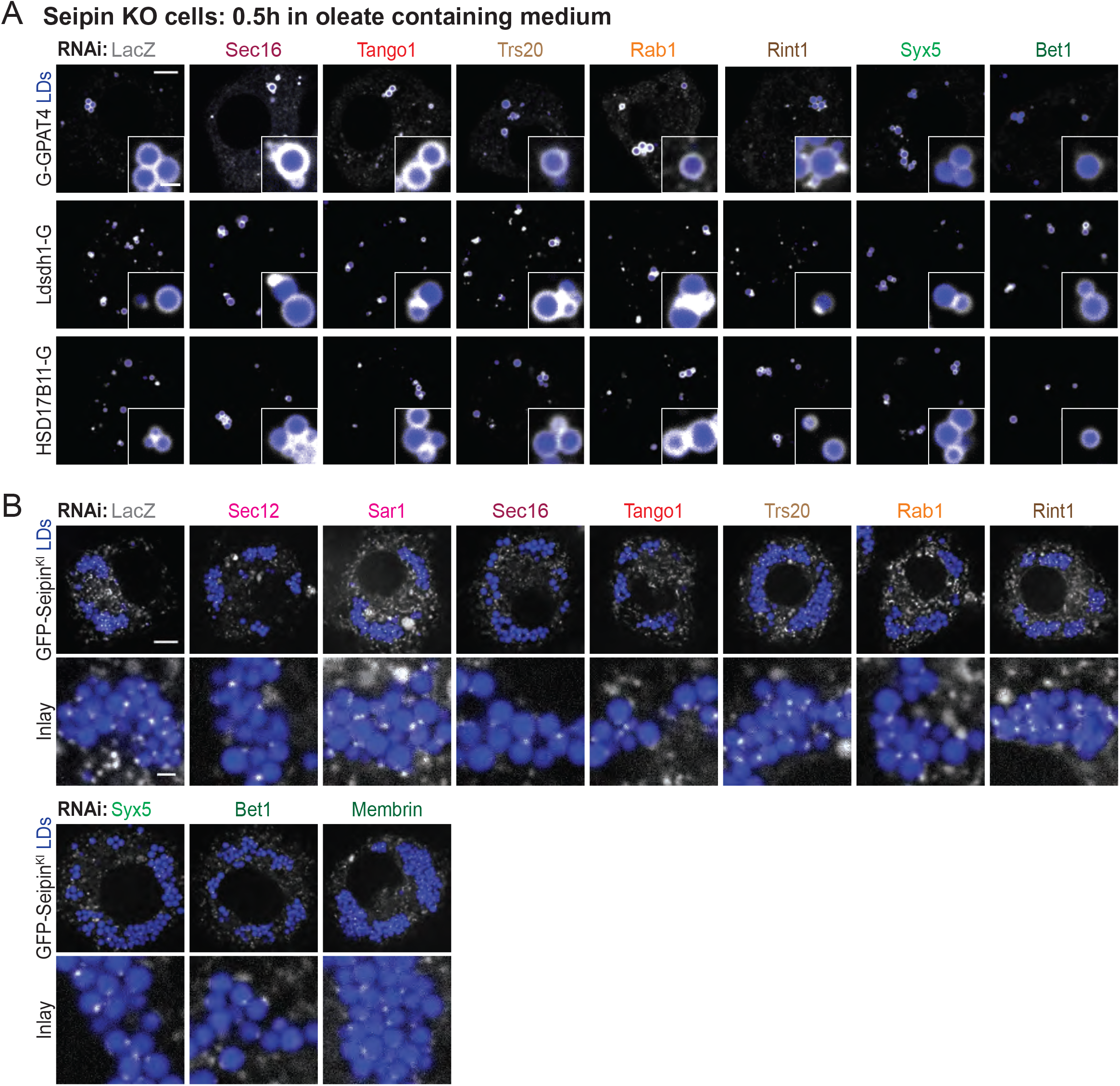
Seipin-mediated ER-LD contacts are independent of late ERTOLD pathway, related to Figure 5. (A) Seipin depletion allows for early targeting of late ERTOLD cargoes even in the absence of late ERTOLD machinery. Confocal imaging of live wildtype or Seipin knock-out cells upon RNAi of ERES or fusion machinery components, followed by transient transfection with EGFP tagged constructs and a 0.5-hr incubation in oleate-containing medium. Scale bar, 5 and 1 μm (inlay). Quantification is shown in Figure 5D. (B) Depletion of late ERTOLD machinery does not affect association of seipin puncta with LDs. Confocal imaging of live GFP-Seipin endogenous knock-in cells upon RNAi of ERES or fusion machinery components, followed by a 20-hr incubation in oleate-containing medium. Scale bar, 5 and 1 μm (inlay).

**Figure S7.**
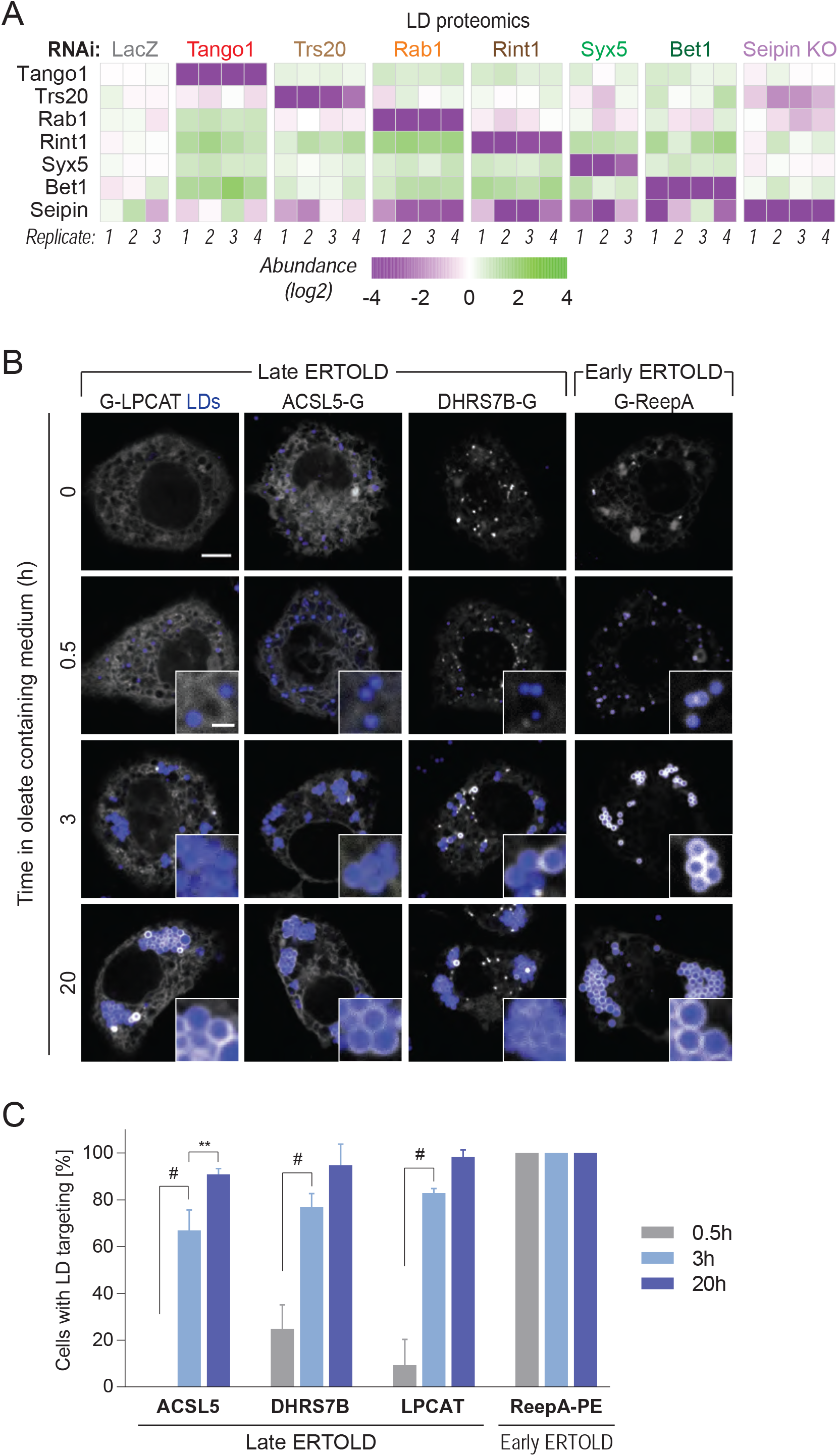
LD proteomics reveal additional late ERTOLD cargoes, related to Figure 6. (B) Heatmap of abundance of ERES and fusion-machinery components and seipin in LD fractions upon their depletion by RNAi (or gene deletion for Seipin) compared to LacZ control, as measured by mass spectrometry. (C) REEPA targets LDs early, and LPCAT, ACSL5, and DHRS7B target LDs late upon LD induction. Confocal imaging of live wildtype cells transiently transfected with EGFP-tagged constructs at given timepoints after 1 mM oleic acid treatment. LDs were stained with MDH. Representative images are shown. Scale bar, 5 and 1 μm (inlay). (D) Bar graph showing percentage of cells with LD targeting over time from the imaging experiment in (B). Mean ± SD, n = 3 independent experiments (10–20 cells each). One-way ANOVA, **p<0.01, #p<0.0001.

## METHODS

- KEY RESOURCES TABLE
- RESOURCE AVAILABILITY

- Lead Contact
- Materials Availability
- Data and Code Availability
- EXPERIMENTAL MODEL AND SUBJECT DETAILS

- Cell lines and cell culture
- METHOD DETAILS

- Special reagents
- Genome-scale RNAi imaging screen
- Plasmid construction
- Transfection
- *In vitro* double-stranded RNA synthesis
- RNA interference
- Generation of a stable cell line
- Generations of cell lines using CRISPR-Cas9
- Immunofluorescence
- Fluorescence microscopy
- Fractionation of cells
- Immunoblotting
- Mass Spectrometry
- QUANTIFICATION AND STATISTICAL ANALYSIS

- Statistical analysis
- Analysis of genome-scale imaging screen
- Quantification of fluorescence images
- Spatial association between ERES and LDs
- Quantification of immunoblots
- Analysis of mass spectrometry data

### KEY RESOURCES TABLE

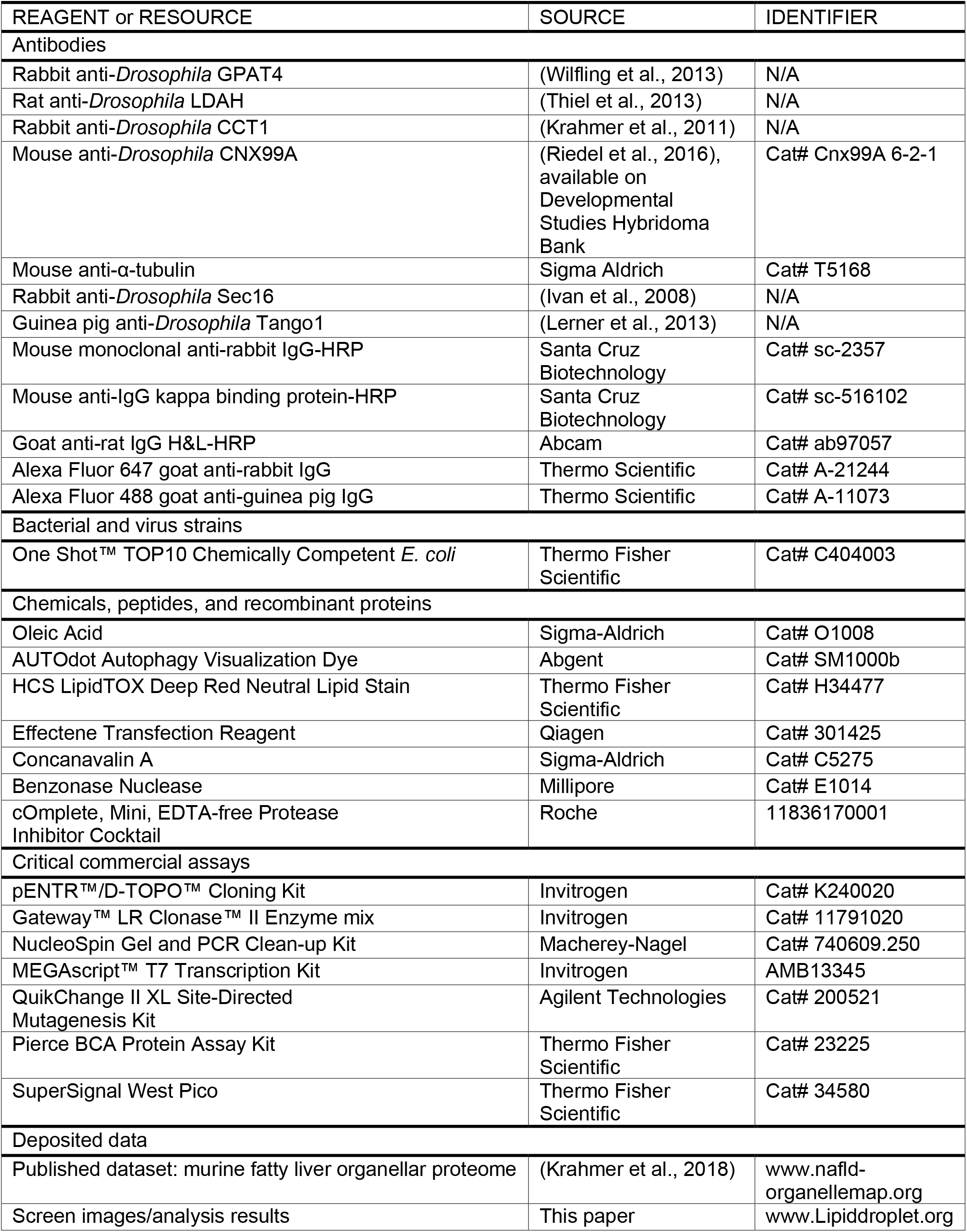
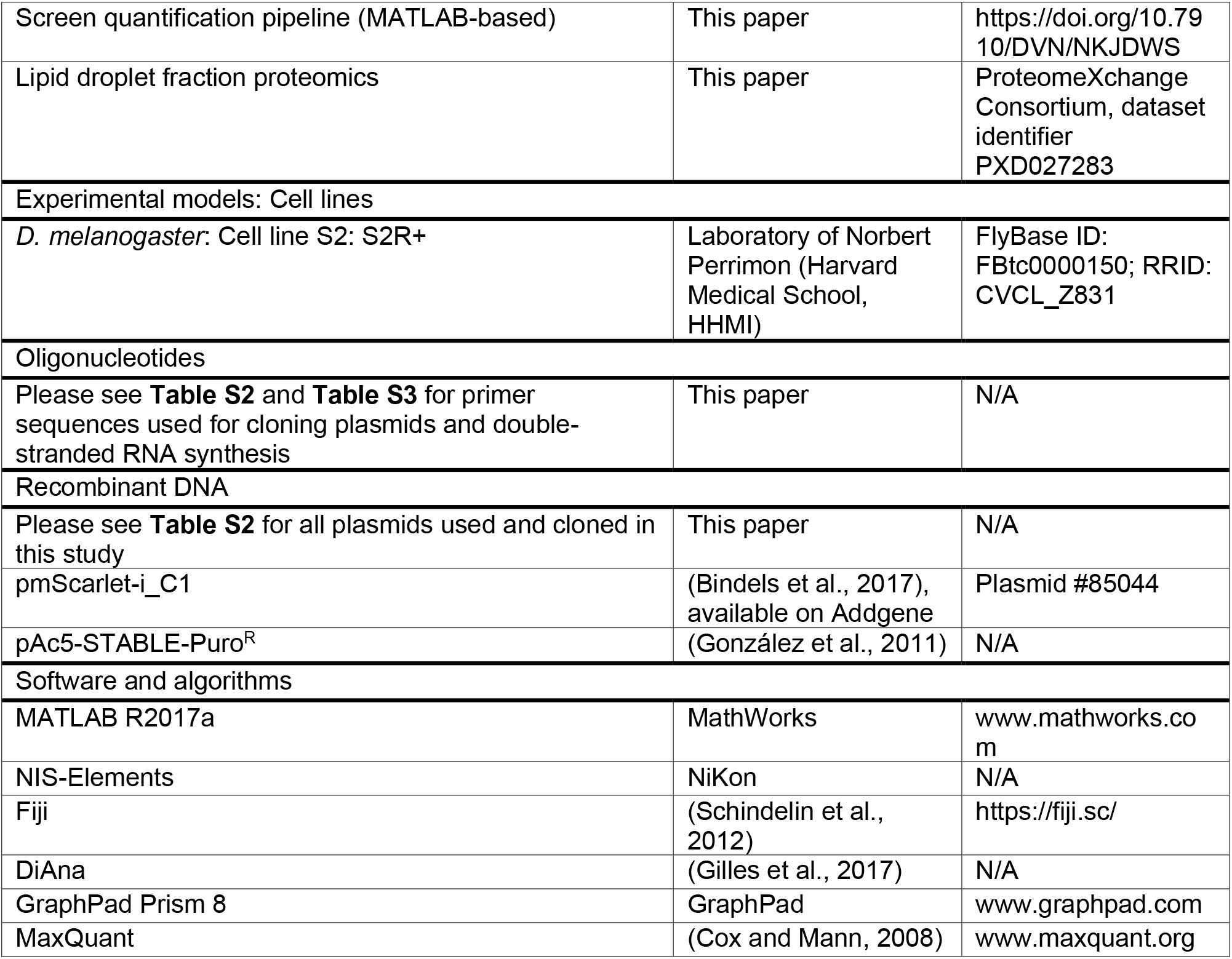

### RESOURCE AVAILABILITY

#### Lead Contact

Further information and requests for resources and reagents should be directed to and will be fulfilled by the Lead Contact, Tobias C. Walther.

#### Materials Availability

All unique/stable reagents used in this study are available from the Lead Contact with a completed Materials Transfer Agreement in accordance with the Harvard T.H. Chan School of Public Health policies.

#### Data and Code Availability

Original screen images and quantification results will be available at the Lipid Droplet Knowledge Portal [http://lipiddroplet.org/; (Mejhert et al., 2021)]. All original code (used for quantification of screen images) has been deposited at Harvard Dataverse and is publicly available. DOIs are listed in the key resources table.

The mass spectrometry proteomics data have been deposited to the ProteomeXchange Consortium via the PRIDE (Perez-Riverol et al., 2019) partner repository with the dataset identifier PXD027283.

### EXPERIMENTAL MODEL and SUBJECT DETAILS

#### Cell lines and cell culture

The *Drosophila* cells used in this study belong to the S2R+ cell line (sex: male) and were provided by Dr. Norbert Perrimon (Harvard Medical School). Cells were cultured at 26°C in Schneider’s *Drosophila* Medium (Gibco, #21720001) supplemented with 10% fetal bovine serum (Gibco), 25 units/ml penicillin, and 25 μg/ml streptomycin. Cells were maintained by splitting 1:6–1:12 every 3–4 days. Where noted, cells were incubated with 1 mM oleic acid complexed with essentially fatty acid–free BSA to induce LDs.

### METHODS DETAILS

#### Special reagents

*Janelia Fluor 646 HaloTag Ligand (JF646)* was a kind gift from Dr. Luke Lavis (Janelia Research Campus, USA). Anti-*dm*LDAH used for western blot experiments (Thiel et al., 2013) was a gift from Dr. Mathias Beller (Heinrich Heine Universitat Dusseldorf, Germany). Anti-*dm*Sec16 used for immunofluorescence experiments (Ivan et al., 2008) was a gift from Dr. Catherine Rabouille (Hubrecht Institute, Netherlands). Anti-*dm*Tango1 used for immunofluorescence experiments (Lerner et al., 2013) was a gift from Dr. Sally Horne-Badovinac (University of Chicago, USA).

10 mM oleic acid was prepared by dissolving 1.98 g of essentially fatty acid–free BSA in 10 mL PBS, adding 31.74 μL oleic acid drop-by-drop, and shaking at 37°C for 1 hr. Solution was sterile-filtered (0.22 μm) before use. All oleic acid treatments were performed with 1 mM final concentration.

### Genome-scale RNAi imaging screen

*Drosophila* S2 R+ cells stably overexpressing EGFP-GPAT4 were subjected to a genome-scale library of dsRNA in imaging-compatible 384-well plates (PerkinElmer, #6057300) two times, prepared by the HMS *Drosophila* RNAi Screening Center (DRSC 2.0 genome-wide screening library). The library targets approximately 13,900 genes 1∼2 times and consists of 66 384-well plates with 250 ng of double-stranded RNA (dsRNA) in 5 μL per well. Confluent cells were resuspended from plates to 60×10^4^ cells/mL in Schneider’s *Drosophila* Medium (Gibco, #21720001) without serum supplementation. 10 μL of the cell suspension was dispensed into the dsRNA plates using the Thermo Scientific Matrix WellMate Microplate Dispenser. After mixing the contents gently, plates were sealed with parafilm and placed in a ‘wet chamber’ (airtight container with wet paper towels) in a 26°C incubator for 50 min. 30 μL of Schneider’s *Drosophila* Medium supplemented with 10% fetal bovine serum, 100 units/mL of penicillin, and 100 μg/mL of streptomycin was added to each well, and plates were sealed with parafilm and placed in the wet chamber for 3.75 days.

After RNAi, 6uL of 10 mM OA solution and 14 μL of fresh media was dispensed to each well, and the plates were sealed with parafilm and placed in the wet chamber for 20 hr. Next day, wells were washed once with 50 μL of PBS and about half the liquid was carefully aspirated using vacuum aspirator to avoid disrupting cells at the bottom of the plate, leaving ∼50 μL. 50 μL of freshly prepared 8% paraformaldehyde in PBS solution was dispensed to each well and incubated at room temperature for 25 min. Again with careful aspiration, wells were washed with 70 μL of PBS three times. 17 μL of 1 μM SiR-DNA nuclear stain (Spirochrome, #SC007), and 133 μM monodansylpentane LD stain (AUTOdot; Abcepta, #SM1000b) in PBS was added to each well (final concentration 0.25 μM SiR-DNA & 33.3 μM AUTOdot) and incubated for 35 min. Finally, each well was washed with 70 μL of PBS three times, and 25 μL of PBS was added (final volume ∼75 μL) for imaging.

For automated confocal imaging, GE IN Cell Analyzer 6000 Cell Imaging System with robotics support for automated plate loading. Using the IN Cell Analyzer software, three channel images (FITC for EGFP-GPAT4, Cy5 for nuclei, and DAPI for LDs) were taken in eight fields per well at the manually determined offset from auto-focusing for each plate using 60X objective.

### Plasmid construction

PCR of the insert was performed using PfuUltra II Fusion Hotstart DNA Polymerase (Agilent Technologies, #600672), following the manufacturer’s protocol. Purified PCR product was first cloned into an entry vector using the pENTR/D-TOPO Cloning Kit (Invitrogen, #K240020) and then into a destination vector from the *Drosophila* Gateway vector collection system (Murphy Laboratory, Carnegie Mellon University) using the Gateway LR clonase Enzyme mix (Invitrogen, #11791019).

Halo destination vectors were created by replacing the EGFP sequence of pAGW and pAWG from the *Drosophila* Gateway vector collection system with Halo sequence using restriction-ligation (EcoRV & AgeI and SacI & AgeI, respectively).

For mutagenesis of constructs, mutations were made using the QuikChange II XL Site-Directed Mutagenesis Kit (Agilent, #200521) in an entry vector, which was then cloned into a destination vector to avoid undetected mutations.

All final plasmids were verified by restriction analysis and sequencing of the insert region. All information about PCR template and primers are provided in **Table S2**.

#### Transfection

Cells were transfected with the Effectene Transfection Reagent (Qiagen, #301425), following the manufacturer’s protocol. When co-transfecting with more than one plasmid, equal amount (in μg) of the plasmids were used. Any further treatments took place 26 hr after transfection.

#### *In vitro* double-stranded RNA synthesis

Genomic DNA of the cells was isolated using the DNeasy Blood & Tissue Kit (Qiagen, #69504). PCR was performed using primers containing the T7 promoter sequence (on both forward and reverse primers) with PfuUltra II Fusion Hotstart DNA Polymerase (Agilent Technologies, #600672). PCR products with the expected size was separated using 1% agarose gel. Purified PCR products were used as a template for RNA transcription using the MEGAscript T7 Transcription Kit (Invitrogen, #AM1334), which was then purified using the RNeasy Mini Kit (Qiagen, #74104). Sizes and quality of synthesized dsRNA were confirmed using 1% agarose gel before they were used for RNAi. PCR primer sequences are provided in **Table S3**.

#### RNA interference

Cells were spun down at 300g for 5 min and resuspended with Schneider’s *Drosophila* Medium (Gibco, #21720001) without serum supplementation at 60×10^4^ cells/mL. Cells were plated first and dsRNA was added at 20 ng/μL. After carefully shaking the plate to mix the contents, plates were sealed with parafilm and placed in a ‘wet chamber’ (airtight container with wet paper towels) inside 26°C incubator to prevent evaporation for 50 min. After 50 min, serum-supplemented medium with three volumes of initial cell suspension were added carefully to avoid detaching cells. After sealing with parafilm, the plate was incubated in the wet chamber for 3.5–4 days before further treatments. In transfection, cells were transferred onto a new plate before following the transfection protocol.

#### Generation of a stable cell line

A stable cell line overexpressing EGFP-GPAT4 was created by transfecting cells with pActin-EGFP-GPAT4-T2A-Puro^R^. Information on PCR template and primers used for cloning the construct is provided in **Table S2**. Selection was started 3 days after transfection with 10 μg/mL puromycin. 5 days later when most control cells (>75%) have died, transfected cells were recovered in medium without puromycin for 5 days. This selection and recovery were repeated one more time before cells were sorted.

For cell sorting, cells were suspended in sterile PBS supplemented with 1% fetal bovine serum. Using FACSAria-561 with 100 μm gating, EGFP+ cells (488 nm laser) were sorted into a 96-well plate (100 cells/well) containing conditioned medium (medium collected from cells growing at exponential phase, combined with equal volume of fresh Schneider’s medium supplemented with 20% fetal bovine serum). After 2 weeks, cells were expanded and subjected to microscopy and western blot for verification of the cell line.

#### Generations of cell lines using CRISPR-Cas9

Knock-out and knock-in cell lines were created using CRISPR-Cas9, following protocols published by Housden et al. (Housden and Perrimon, 2016; Housden et al., 2016). Guide RNA sequence and PCR primer sequences for donor construct cloning are provided in **Table S2**. 1 week after transfection, cells were suspended in sterile PBS supplemented with 1% fetal bovine serum and single-cell sorted into a 96-well plate containing conditioned medium (medium collected from cells growing at exponential phase, combined with equal volume of fresh Schneider’s medium supplemented with 20% fetal bovine serum), using FACSAria-561 with 100-μm gating. After 2∼3 weeks, viable single-cell colonies were expanded and subjected to microscopy, western blot, and sequencing for verification of correct genome-editing.

#### Immunofluorescence

Cells were plated to a 96-square well clear bottom plates (Perkin Elmer) in 1:1 dilution with fresh media (100 μL of 80∼90% confluent cell suspension + 100 μL of fresh medium). After 1 hr, 22.2 μL of 10 mM oleic acid was added. At a given OA timepoint, 100 μL of liquid was removed (leaving ∼100 μL, careful not to dry the well or disrupt the cells at the bottom of the well). Wells were washed 2 times with 200 μL of PBS. 200 μL of 6% paraformaldehyde (Polysciences 18814-10) in PBS was added to each well and incubated for 10 min, followed by washing with 200 μL of PBS 4 times. Cells were permeabilized with 200 μL of 0.15% triton X-100 and 0.15% BSA in PBS for 3 min, washed 4 times with 200 μL of PBS, blocked with 200 μL of 7.5% normal goat serum (Cell Signaling 5425S) in PBS for 1 hr. After removing 200 μL from the well (leaving ∼100 μL), 50 μL of antibody solution for the final concentration of 1:1500 rabbit anti-*dm*Sec16 (Ivan et al., 2008) or guinea pig anti-*dm*Tango1 (Lerner et al., 2013) in 5% normal goat serum in PBS was added and incubated for 1.5 hr. After removing 50 μL from the well (leaving ∼100 μL), and wells were washed with 0.2% BSA in PBS solution 4 times with incubation for 5 min with each wash. Wells were washed once with 200 μL of 5% normal goat serum in PBS. 50 μL of secondary antibody solution for the final concentration of 1:1000 (Alexa Fluor 488 goat anti-guinea pig IgG, Thermo Scientific A-11073; Alexa Fluor 647 goat anti-rabbit IgG, Thermo Scientific A-21244) in 5% normal goat serum in PBS was added and incubated for 1.5 hr. After removing 50 μL from the well (leaving ∼100 μL), wells were washed with 0.2% BSA in PBS solution 4 times with incubation for 5 min with each wash, followed by washing with 200 μL of PBS 4 times. Finally, 0.10 μL AutoDot (Abcepta SM1000b) in 200 μL of PBS was added for LD staining before imaging.

#### Fluorescence microscopy

Cells that have undergone transfection or RNAi in 24-well plates were resuspended in the old medium and combined with equal volume of fresh medium to a 35-mm dish with 14-mm No. 1.5 coverslip bottom (MatTek Life Sciences, #P35G-1.5-14-C) coated manually with 0.1 mg/mL Concanavalin A. Cells were allowed to settle for 1 hr at 26°C before further treatments, such as with 1 mM oleic acid (OA). Unless otherwise indicated, cells were imaged 20 hr after OA treatment. LDs were stained with 100 μM monodansylpentane (AUTOdot; Abcepta, #SM1000b) or 1:1000 HCS LipidTOX Deep Red Neutral Lipid Stain (Thermo Fisher Scientific, H34477), 10 min before imaging. For JF646 treatment, cells were incubated with the Halo ligand 1 hr before imaging and washed once with PBS before resupplying medium (or medium with OA).

Nikon Eclipse Ti inverted microscope, featuring CSU-X1 spinning disk confocal (Yokogama) and Zyla 4.2 PLUS scientific complementary metal-oxide semiconductor (sCMOS) (Andor, UK), was used for spinning disk confocal microscopy. NIS-elements software (Nikon) was used for acquisition control. Plan Apochromat VC 100X oil objective (Nikon) with 1.40 NA was used, resulting in 0.065-μm pixel size. Solid state excitation lasers—405 nm (blue; Andor), 488 nm (green; Andor), 560 nm (red; Cobolt), and 637 nm (far-red; Coherent)—shared quad-pass dichroic beam splitter (Di01-T405/488/568/647, Semrock), whereas emission filters were FF01-452/45, FF03-525/50, FF01-607/36, and FF02-685/40 (Semrock), respectively.

#### Fractionation of cells

Cells were resuspended from plates and washed once with PBS, which was then placed on ice for all subsequent steps. Cell pellets were suspended in 1mL of 250 mM sucrose buffer containing 200 mM Tris-HCl (pH 7.4), 1 mM MgCl_2_ (pH 7.4), and cOmplete Mini EDTA-protease inhibitor cocktail (Roche, #4693159001) and broken by passing through 25G syringe 30 times. 1unit/μL of benzonase nuclease (Millipore, #E1014) was added and incubated on ice for 10 min. 5% of the total volume was taken at this stage for whole-cell lysate (‘input’) analysis. For the rest, unbroken cells and nuclei were fractionated by centrifuging for 5 min, 1,000g at 4°C. Top lipid layer and the supernatant were moved to the 5-mL, Open-Top Thinwall Ultra-Clear Tube, 13×51mm (Beckman Coulter, #344057), where additional 1.5 mL of the 250 mM sucrose buffer was added. 2.5 mL of 50 mM sucrose containing 200 mM Tris-HCl (pH 7.4), 1 mM MgCl_2_ (pH 7.4), and cOmplete Mini EDTA-protease inhibitor cocktail was layered on top. The two-step sucrose gradient was centrifuged for 16-20 hr, 100,000g, at 4°C using the SW 55 Ti Swinging Bucket rotor (Beckman Coulter, 342194).

Top of the tube (∼5 mm; 500 μL) was sliced using the Beckman Coulter tube slicer, the content of which was taken as ‘LD fraction’. Supernatant was taken as ‘soluble fraction’, and the pellet resuspended in 500 μL of 250 mM sucrose buffer was taken as ‘membrane fraction’.

For further analysis with immunoblotting or mass spectrometry, proteins from the fractions were precipitated. Briefly, 1 mL of methanol and 250 μL of chloroform was sequentially added to 500∼750 μL of aqueous fractions with vigorous mixing after every addition. After centrifuging for 10 min, 14,000g, at 4°C, top layer was aspirated out. 1.7 mL of methanol was added and vigorously mixed. Precipitated proteins were then pelleted by centrifuging for 15 min, 18,000g, at 4°C. All liquid was aspirated, and after allowing the pellet to dry for 5 min, the pellet was resuspended in 100-250 μL of 1.5% SDS, 50 mM Tris-HCl (pH 7.4) buffer.

#### Immunoblotting

Protein concentrations were measured using the Pierce BCA Protein Assay Kit (Thermo scientific, #23225), and the amounts indicated in respective figure legends were mixed with 5X Laemmli buffer for the final concentration of 1X Laemmli buffer (2% SDS, 10% glycerol, 50 mM Tris-HCl (pH 6.8), β-mercaptoethanol 100 mM, 0.02% bromophenol-blue). After running samples in 4-15% gradient polyacrylamide gel (Bio-Rad, #4561084) at 100V for 90 min in 1X Tris/glycine/SDS buffer (Bio-Rad, #161-0772), proteins were transferred to a 0.2-μm pore size nitrocellulose membrane (Bio-Rad, #1620112) in 1X Tris/glycine buffer (Bio-Rad, #161-0771) at 70V for 90 min in the cold room (4°C). Membrane was blocked by incubating in 5% non-fat dry milk (Santa Cruz Biotechnology, #sc-2325) in TBS-T buffer (20 mM Tris, pH 7.6, 150 mM NaCl, 0.1% Tween-20) for 30 min at room temperature. Membrane was incubated with 5% milk solution containing primary antibody (dilutions are indicated below) overnight in the cold room.

Next day, membranes are washed three times with TBS-T for 10 min each at room temperature, incubated with 1:5000 secondary antibodies conjugated to horseradish peroxidase (Santa Cruz Biotechnology, #sc-516102 for primary antibodies from mouse or #sc-2357 for primary antibodies from rabbit) in 5% milk solution for 1 h at room temperature, and washed three times with TBS-T for 10 min each at room temperature. SuperSignal West Pico PLUS Chemiluminescent Substrate (Thermo Scientific, #34580) was applied to the membrane, which was imaged using the Biorad Gel Doc XR system for signal acquisition.

For stripping the membrane of antibodies, membrane was washed with distilled water five times for 5 min each at room temperature and incubated with 100 mM citric acid solution in distilled water for 10 min at room temperature. The membrane was then re-blocked with 5% milk solution for 30 min at room temperature before proceeding with incubation with another primary antibody.

Primary antibodies and their dilutions: rabbit anti-*dm*GPAT4 [1:1000; (Wilfling et al., 2013)], rabbit anti-*dm*CCT1 [1:1000; (Krahmer et al., 2011)], mouse anti-*dm*CNX99A [1:500; DSHB, #Cnx99A 6-2-1, (Riedel et al., 2016)], mouse anti-α-tubulin (1:2000; Sigma, T5168), and anti-*dm*LDAH [1:2000; (Thiel et al., 2013)].

#### Mass spectrometry

Proteins pellets from lipid droplet-enriched fractions were resuspended in 0.1 M NaOH (Sigma-Aldrich) and subsequently neutralized using 200 mM HEPES (4-(2-hydroxyethyl)-1-piperazineethanesulfonic acid). Solubilized proteins were reduced using 5 mM dithiothreitol (Sigma-Aldrich), pH 7.5, at 37°C for 1 hr. Reduced disulfide bonds of cysteine residues were alkylated using 15 mM iodoacetamide (Sigma-Aldrich) for 1 hr in the dark. Excessive iodoacetamide was quenched using 10 mM dithiothreitol. Alkylated protein mixture was diluted sixfold (v/v) using 20 mM HEPES, pH 7.5, and digested for 16 hr at 37°C with sequencing grade trypsin (Worthington Biochemical) in a 1:100 trypsin to protein ratio. Digested peptides were desalted using self-packed C18 STAGE tips (3M Empore^TM^) (Rappsilber et al., 2003). Desalted peptides were dissolved in 0.1% (v/v) formic acid and injected onto an Easy-nLC 1000 (Thermo Fisher Scientific), coupled to an Orbitrap Exploris 480 (Thermo Fisher Scientific). Peptide separation was performed on a 500-mm self-packed analytical column using PicoTip^TM^ emitter (New Objective, Woburn, MA) containing Reprosil Gold 120 C-18, 1.9-µm particle size resin (Dr. Maisch, Ammerbuch-Entringen, Germany). Chromatography separation was carried out using increasing organic proportion of acetonitrile (5–40 % (v/v)) containing 0.1 % (v/v) formic acid over a 120 min gradient at a flow rate of 300 nL/min.

### QUANTIFICATION AND STATISTICAL ANALYSIS

#### Statistical analysis

All statistical analysis was performed using GraphPad Prism 8. Information about significance test is provided in the respective figure legends. All multiple comparisons were performed with the Bonferroni multiple comparisons correction. Statistically significant differences are denoted as follows: *p<0.05, **p<0.01, ***p<0.001, #p<0.0001.

#### Analysis of genome-scale imaging screen

A custom Matlab analysis pipeline was built to analyze screen images. Nuclei, cell, and LD compartments were obtained using supervised machine-learning methods, specifically Random Forest pixel classifiers (http://github.com/HMS-IDAC/PixelClassifier). Three different models were trained, one for each compartment, using separate sets of annotated images (about seven images per model). Nuclei and cells were segmented from the Cy5 channel, and LDs formed the DAPI channel. Nuclei mask was used for segmenting cells as markers in a watershed algorithm. LD masks were post-processed by selecting only the intersection with cell masks. LD objects were then associated with cell objects, depending on the area of intersection. Finally, the signal in the FITC channel was corrected for auto-fluorescence by subtracting the mean value of control images and was quantified inside and outside the LD mask in each segmented cell. From these measurements, we calculated LD targeting ratio for each segmented cell, defined as the ratio of the mean intensity of EGFP-GPAT4 signal inside the LD mask to that of EGFP-GPAT4 signal outside LD mask within the cell mask. Median LD targeting ratio from all of the segmented cells from the 8 fields of the same well (with a unique dsRNA) was determined and employed as the final readout for the well. Batch patch processing of screen images (total 1,216,512 images) through the custom analysis pipeline was performed using the Harvard O2 cluster.

For the calculation of robust Z-scores, each median LD targeting ratio from the duplicate screen experiment was considered as a separate value. Robust Z-scores for median LD targeting ratios (X) was calculated using the formula below. In our screen, median = 2.147287 and median absolute deviation = 0.113917.

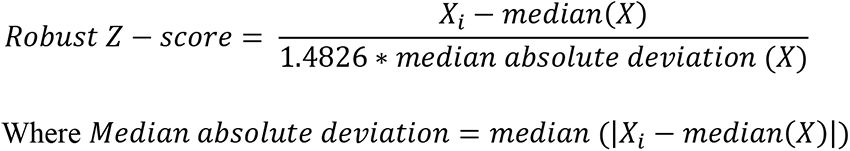

#### Quantification of fluorescence images

Confocal images were quantified using the FIJI software (Schindelin et al., 2012) to calculate LD targeting ratios. Cell boundaries were drawn manually based on fluorescence from protein channels, such as EGFP-GPAT4 (Mask 1) and LD regions, were distinguished by applying an automatic threshold (Otsu method) to the monodansylpentane channel within Mask 1, which was then dilated by 1 pixel to include LD surfaces (Mask 2). LD targeting ratios were calculated by dividing the mean intensity of the fluorescent protein channel image in Mask 2 divided by that in (Mask 1 – Mask 2). For CCT1, nuclei were distinguished by manually drawing the nuclear boundary from mCherry-CCT1 channel (Mask 3), which was excluded from Mask 1.

#### Spatial association between ERES and LDs

The ERES-LD association was determined by DiAna, an ImageJ plugin for object-based 3D co-localization analysis (Gilles et al., 2017). The fluorescence signal of ERES punctate (Sec16 or Tango1) and LDs were segmented by intensity thresholding. The intensity threshold of the segmentation was automatically determined by an algorithm. After the thresholding, the ERES and LD were identified as 3D objects, and the number of ERES and the closest distance between the ERES and LD boundaries were determined.

The images were preprocessed by background subtraction and median filter before the segmentation. To exclude the interference from cell debris and non-specific labelling, only the objects with pixel size > 30 pixels were considered in the analysis. The objects located at the x, y and z boundaries of the image stack were also excluded from the calculation to prevent the error from the incomplete segmentation of the objects. Zero closest distance between ERES and LDs indicated overlapping boundaries between the two. The ratio of the ERES associated with LD was calculated by dividing the number of ERES with zero closest distance with LDs by the total number of ERES within the cell.

#### Quantification of immunoblots

Using the FIJI software (Schindelin et al., 2012), a rectangular ROI was drawn around the band of interest in the control lane (usually LacZ RNAi), and the measure function was used to measure total intensity inside the ROI. ROI was then moved to the next lane sequentially for measurements. Another measurement outside the expected band region was measured for background subtraction. The measured intensities (band – background) were normalized to the level in the control lane.

#### Analysis of mass spectrometry data

The mass spectrometry analyzer operated in data-dependent acquisition mode with a top 10 method at a mass-over-charge (m/z) range of 300-2000 Da. Mass spectrometry data were analyzed by MaxQuant software version 1.5.2.8 (Cox and Mann, 2008) using the following setting: oxidized methionine residues and protein N-terminal acetylation as variable modification, cysteine carbamidomethylation as fixed modification, first search peptide tolerance 20 ppm, main search peptide tolerance 4.5 ppm. Protease specificity was set to trypsin with up to 2 missed cleavages allowed. Only peptides longer than 6 amino acids were analyzed, and the minimal ratio count to quantify a protein is 2. The false discovery rate was set to 5% for peptide and protein identifications. Database searches were performed using the Andromeda search engine integrated into the MaxQuant software (Cox et al., 2011) against the UniProt *Drosophila melanogaster* database containing 20,981 entries (December 2018). “Matching between runs” algorithm with a time window of 0.7 min was employed to transfer identifications between samples processed using the same nanospray conditions. Protein tables were filtered to eliminate identifications from the reverse database and common contaminants.

To identify proteins regulated by different genotypes, the MaxQuant output files were exported to Perseus 1.5.1.6 (Tyanova et al., 2016). Known contaminant and decoy sequences were removed. Projection and clustering of the dataset was performed using principal component analysis to identify potential sample outlier. The cutoff of potential principal components was set at Benjamini-Hochberg FDR 5%. After removing poorly clustering replicates in the principal component analysis, intensity values were normalized to the sum of all intensities within each sample. Clustergram analysis was performed using the ComplexHeatmap clustering method on R (Gu et al., 2016).

### SUPPLEMENTAL TABLE TITLES and LEGENDS

**Supplemental Table 1.** Gene nomenclature used in the study

List of gene and protein names used in this study. For some fly genes and proteins without common names besides an annotation symbol, names of their predicted human orthologs (based on FlyBase.org and Marrvel.org) were used instead. For ortholog predictions, comma ‘,’ denotes aliases for the same protein and semi-colon ‘;’ denotes more than one possible ortholog. For ReepA, ReepA-RE isoform sequence used in the study for expression.

**Supplemental Table 2.** Plasmid construction for transient transfection or cell line generation

List of primer sequences, PCR template, and backbone information for the construction of plasmids used for transient transfection (first tab) or for the generation of cell lines (second tab) used in the study.

**Supplemental Table 3.** Oligonucleotide sequences used for the synthesis of double-stranded RNA

List of primer sequences for generating PCR templates used in *in vitro* transcription. All primers were flanked by T7 promoter sequence (TAATACGACTCACTATAGGG) at the 5′ end.

