## Supplemental Table 1 for "Identification of two pathways mediating protein targeting from ER to lipid droplets"

**Table S1**. Gene nomenclature used in the study

| **Notation used in the paper** | **FlyBase ID** | **Annotation Symbol** | **Fly common name** | **Human ortholog name(s)** |
| --- | --- | --- | --- | --- |
| ***ER-LD Bridge Factors*** | | | | |
| Sec12 | FBgn0031779 | CG9175 | CG9175 | PREB, **Sec12** |
| Sar1 | FBgn0038947 | CG7073 | **Sar1** | Sar1b; Sar1a |
| Sec16 | FBgn0052654 | CG32654 | **Sec16** | Sec16A; Sec16B |
| Sec23 | FBgn0262125 | CG1250 | **Sec23** | Sec23A; Sec23B |
| Tango1 | FBgn0286898 | CG11098 | **Tango1** | MIA2; CTAGE8 |
| Trs20 | FBgn0266724 | CG5161 | **Trs20** | TRAPPC2, sedlin |
| Rab1 | FBgn0285937 | CG3320 | **Rab1** | Rab1A; Rab1B |
| Rint1 | FBgn0035762 | CG8605 | **Rint1** | Rint1 |
| Syx5 | FBgn0011708 | CG4214 | **Syx5** | STX5 |
| Membrin | FBgn0260856 | CG4780 | **Membrin** | GOSR2 |
| Bet1 | FBgn0260857 | CG14084 | **Bet1** | Bet1; Bet1L |
| Ykt6 | FBgn0260858 | CG1515 | **Ykt6** | Ykt6 |
| NSF | FBgn0000346 | CG1618 | comt | **NSF** |
| αSNAP | FBgn0250791 | CG6625 | **αSNAP** | NAPA; NAPB |
| ***LD targeting proteins*** | | | | |
| LDAH | FBgn0035206 | CG9186 | sturkopf | **LDAH** |
| Ubxd8 | FBgn0025608 | CG10372 | Faf2 | Faf2, **Ubxd8** |
| GPAT4 | FBgn0034971 | CG3209 | **GPAT4** | GPAT4; GPAT3 |
| Ldsdh1 | FBgn0029994 | CG2254 | **Ldsdh1** | HSD17B11 |
| HSD17B11 | FBgn0032910 | CG9265 | CG9265 | SDR16C5; **HSD17B11** |
| LPCAT | FBgn0052699 | CG32699 | **LPCAT** | LPCAT1; LPCAT2 |
| ACSL5 | FBgn0036821 | CG3961 | CG3961 | **ACSL5**; ACSL1 |
| DHRS7B | FBgn0027583 | CG7601 | CG7601 | **DHRS7B** |
| REEPA | FBgn0261564 | CG42678 | **ReepA** | REEP2; REEP1; REEP4 |
| Lsd1 | FBgn0039114 | CG10374 | **Lsd1** | PLIN2; PLIN3 |
| CGI-58 | FBgn0033226 | CG1882 | puml | ABHD4; ABHD5, **CGI-58** |
| CCT1 | FBgn0041342 | CG1049 | Pcyt1, **CCT1** | PCYT1A; PCYT1B |

List of gene and protein names used in this study. For some fly genes and proteins without common names besides an annotation symbol, names of their predicted human orthologs (based on FlyBase.org and Marrvel.org) were used instead. For ortholog predictions, comma ',' denotes aliases for the same protein and semi-colon ';' denotes more than one possible ortholog. For ReepA, ReepA-RE isoform sequence used in the study for expression.
